# Dietary methionine restriction primes T cell metabolism for activation and tumor inhibition and enhances the efficacy of immune checkpoint blockade

**DOI:** 10.1101/2025.09.18.676961

**Authors:** Xiaoqing Qing, Binita Chakraborty, Panpan Liu, Sandeep Artham, Patrick K Juras, Dongyin Guan, Kristen E. Pauken, Ching-Yi Chang, Xia Gao

## Abstract

The proliferation of many cancer cells is methionine dependent and dietary methionine restriction (MR) has shown anti-tumor effects in a wide variety of immunodeficiency preclinical models. Yet, whether MR exerts an anti-tumor effect in the presence of an immune-competent background remains inconclusive. Accumulating evidence has shown an essential role of methionine in immune cell differentiation and function. Thus, competition for methionine between tumor cells and immune cells in the tumor microenvironment may drive tumor growth and tumor response to therapy. Here, we aim to define the impact of MR on tumor growth and associated immunity. We first assessed the effect of MR in a series of immunocompetent mouse models of melanoma, colorectal cancer, breast cancer, and lung. MR led to a broad tumor inhibition effect across these models and such tumor inhibition was not sex-or genetic background-dependent but appears to be fully or partially immune-dependent. Through flow cytometry analysis, we found a consistent increase in intratumoral activated CD8^+^ T cells across different tumor models and depletion of CD8^+^ T cells partially or completely reversed MR-induced tumor inhibition in a model dependent manner. Interestingly in young healthy non-tumor-bearing mice, MR increased spleen CD3^+^ and CD8^+^ T cell populations. Metabolomics and RNAseq analysis of spleen-derived CD8^+^ T cells revealed significant increase in purine metabolism and amino acid metabolism and that are in line with the metabolic feature of activated T cells. Furthermore, MR improved the efficacy of anti-PD1 immune checkpoint blockade. Together, MR primes T cell metabolism for its anti-tumor effect and improves the efficacy of anti-PD1 checkpoint blockade.

## Introduction

Diet affects every aspect of cancer, from initiation to development and to the response to therapy. It has been estimated that diet contributes to ∼20% of cancer-related mortality, and ∼ one-third of most common cancers are preventable through lifestyle change, including changing the diet^1–3^. Thus, dietary intervention holds a strong promise for cancer prevention and therapy^2,4–6^.

One intriguing possibility of dietary intervention for cancer therapy is the restriction of methionine^7^. Methionine, an essential sulfur-containing amino acid, is involved in a myriad of cellular functions: biosynthesis, redox balance, epigenetic stability, and signaling. It is highly variable in human circulation that can be manipulated by dietary intervention^7,8^. Dietary methionine restriction (MR) has shown numerous health benefits including anti-aging^9,10^, anti-obesity and associated metabolic syndromes^11–13^, anti-inflammation^14–16^, and anti-tumor^17^. Unlike normal healthy cells, many cancer cells exhibit different extents of methionine auxotrophy^18^. Mounting evidence supporting an anti-tumor role of MR^7,19–24^.

While methionine is essential for cancer cells, methionine-centered one-carbon metabolism is also indispensable for T cell activation and function^14,25,26^. CD4^+^ and CD8^+^ T cell activation is accompanied by significant uptake of methionine^26,27^ and dramatic increase in S-adenosyl methionine (SAM), which mediates cell differentiation program through histone methylations^14,25,26^. The primary source of methionine for T cells is the exogenous supply^14^.

Since both tumor cells and T cells require methionine for their growth and function, it is possible that tumor cells and T cells may compete for methionine in the tumor microenvironment (TME). Indeed, recent studies suggested tumor cells in the TME could outcompete T cells for methionine uptake via transporter SLC43A2, resulting in T cell exhaustion via reduced methionine availability, SAM and SAM-mediated H3K79me2 histone methylation^28,29^. In contrast to all the other reports^19–21,30^, a recent study by Ji et al.^31^ has further challenged the tumor-suppressing effect of MR in immune-competent backgrounds.

Considering this controversy on the anti-tumor role of MR in immune-competent backgrounds, we aim to determine the relevance of immune system in MR-mediated tumor outcomes. Here, using a series of mouse models for melanoma, colorectal cancer, breast cancer, and lung cancer, we first assessed the impact of MR on tumor growth in the presence of an immune-competent background, and further defined the relevant intratumor immune cells, and the potential mechanisms.

## Results

### MR inhibits tumor growth in immune-competent mouse cancer models

Recent reports have questioned the anti-tumor effect of dietary MR in immune-competent backgrounds^28,29,31^. To provide a definite answer to whether dietary MR has an anti-tumor effect in an immune-competent background, we first conducted dietary intervention studies by feeding mice the control (0.86% methionine) or MR (0.17% methionine) diet in a series of syngeneic mouse models for melanoma, colorectal, and breast cancer. In melanoma syngeneic mouse models, 7-8-week-old C57BL/6J mice, both female and male, were inoculated with BPD6 cells that carry BRAF^V600E^ mutation and PTEN deletion or YUMM5.2 cells that carry BRAF^V600E^ mutation and P53 deletion. In female mice, both BPD6 and YUMM5.2 tumors progressed slower when fed the MR diet compared to mice fed the control diet (**Figure 1A**). Similarly in male mice, MR led to a trend toward tumor inhibition in BPD6 model and a significant tumor inhibition in YUMM5.2 tumors (**Figure 1B**). Additionally, we assessed the effect of MR in a clinically relevant autochthonous melanoma mouse model iBP^32^, in which tumor initiation and growth were driven by concomitant conditional activation of *Braf*^V600E^ and homozygous deletion of *Pten* in melanocytes (Braf^tm1Mmcm^ *Pten^fl/fl^* mTyr-CreERT2). This genetic mouse model resembles human melanomas bearing BRAF mutation and PTEN deletion, similar to the BPD6 syngeneic model. Consistent with the syngeneic melanoma models, MR also significantly inhibited tumor growth in iBP model (**Figure S1A**). Together, MR alone significantly inhibited melanoma tumor growth in two syngeneic models and one genetic mouse model, and such tumor inhibition is not sex-dependent, at least in this age group of mice.

**Fig 1.**
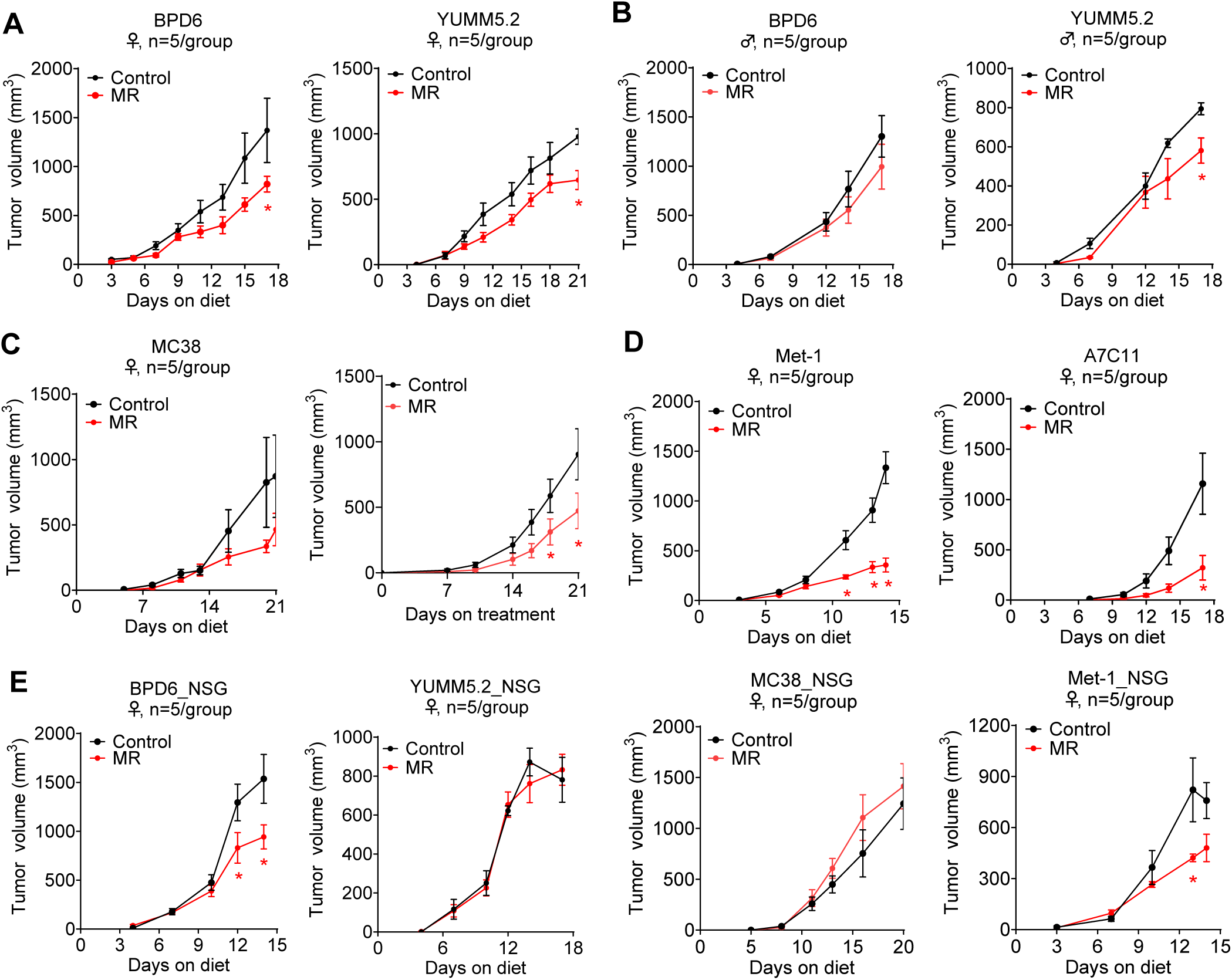
MR inhibits tumor growth in immune-competent mouse cancer models. A. Tumor growth in BPD6 and YUMM5.2 syngeneic melanoma models established in female C57BL/6J mice (*n =* 5/group). B. Tumor growth in BPD6 and YUMM5.2 syngeneic melanoma models established in male C57BL/6J mice (*n =* 5/group). C. Tumor growth in MC38 syngeneic colorectal cancer models established in female (*n =* 5/group) and male (*n =* 8-9/group) C57BL/6J mice. D. Tumor growth in Met-1 and A7C11 syngeneic melanoma models established in female FVB and C57BL/6J mice, respectively (*n =* 5/group). E. Tumor growth in BPD6, YUMM5.2, MC38, and Met-1 models established in female NSG mice (*n =* 5/group). Data were expressed as mean ± SEM. *P *<* 0.05 by 2-way ANOVA followed by Bonferroni’s multiple-correction test.

Since MR showed tumor inhibition in melanoma models, we extended our assessment of MR effect in other syngeneic models of colorectal, breast, and lung cancers. For colorectal cancer models, we used MC38, carrying mutations in P53 and PTEN, established in C57BL/6J mice. Again, MR led to a significant (male) or a trend (female) towards tumor growth inhibition in MC38 model (**Figure 1C**). For breast cancer models, we established two triple-negative breast cancer models, A7C11 and Met-1, in two different mouse strains, C57BL/6J mice and FVB mice, respectively (**Figure 1D**). Consistently, MR led to a strong tumor inhibition in both A7C11 and Met-1 models. Similarly, MR also inhibited tumor growth in the LLC1 hypermutated non-small lung cancer model in female mice (**Figure S1B**). Thus, MR showed a broad tumor inhibition effect in a wide range of cancer models.

Notably, MR resulted in no significant body weight change in both male and female YUMM5.2 models, and female MC38 models, and a minor weight loss compared to normal diet in BPD6, male MC38, A7C11, Met-1, iBP, and LLC1 models (**Figure S1**). Such minor weight loss appears to be model dependent and unlikely cause harm to host metabolic health, as the MR diet containing 0.17% methionine has been shown to reduce body weight gain and fat mass in healthy young mice without impairing growth or lean mass^33^. Altogether, MR inhibits tumor growth in a variety of immune-competent mouse tumor models that are independent of sex, genetic mutations, or mouse strains.

### MR mediated tumor inhibition is immune relevant

To determine the mechanisms related to MR-mediated tumor inhibition, we first performed a bulk RNA sequencing (RNAseq) analysis on collected MC38 and BPD6 tumors from the above animal studies in female mice. In BPD6 tumors (**Figure S2A**), only 102 genes were significantly changed (P value < 0.05, |log_2_FC| > 0.58) by MR, with 85 genes upregulated and 17 genes downregulated. In MC38 tumors (**Figure S2B**), 650 genes were significantly changed (P value < 0.05, |log_2_FC| > 0.58) by MR, with 454 genes upregulated and 196 genes downregulated. GO term analysis of these significantly changed genes showed the top impacted pathways are largely immune relevant among the upregulated genes in both BPD6 and MC38 models. In comparison, no striking consistency was observed in the GO term analysis of the downregulated genes in the two models. Interestingly, among the significantly changed genes, 20 out of 102 in BPD6 tumors and 91 out of 605 in MC38 tumors are related to immune response (**Figure S2C-D**). Two genes are commonly increased in both models, Clec4d (Cd368) and H2-T23, both of which contribute to antigen presenting indirectly or directly^34–36^. Overall, MR leads to significant changes in immune response at a transcriptional level.

To further determine whether MR-mediated tumor inhibition depends on the immune system, we assessed the effect of MR on tumor growth in NOD.Cg-PrkdcScid Il2rgtm1Wjl/SzJ (NSG) mice that lack functional B, T, and NK cells. For this set of study, we randomly examined 4 tumor models, two melanomas (BPD6 and YUMM5.2), one colorectal cancer MC38, and one breast cancer Met-1. In the immune-compromised NSG mice, MR-mediated tumor inhibition was completely abolished in YUMM5.2 and MC38 models, suggesting the anti-tumor effect of MR in these two models depends on a competent immune system (**Figure 1E**). In BPD6 and Met-1 models, MR-mediated tumor inhibition in NSG mice appeared to be similar or less pronounced when compared to that in their corresponding immune-competent background (**Figure 1A and 1D-E**), indicating an immune-independent mechanism such as a tumor autonomous mechanism is in play, along with a possible partial immune-dependency. Thus, the extent of immune-dependency in MR-associated tumor inhibition is model-specific.

### MR activates tumor infiltrated CD8^+^ T cells

Considering the relevance of immune system to MR-mediated tumor inhibition, we then set to determine which immune cells are responsible. We isolated single cells from tumors at the endpoint of the above animal studies, three melanomas (two syngeneic models, BPD6 and YUMM5.2, and one genetically modified model iBP) and one colorectal cancer model (MC38), and performed immune staining followed by flow cytometry analysis. We first assessed myeloid cell populations via probing against CD11b, CD24, CD11c, CD64, F4/80, MHCII, and CD206 markers (**Figure S3A**). There were no common, significant changes in overall myeloid cells (CD11b^+^), monocytes (CD11b^+^Ly6C^+^Ly6G^-^), neutrophils (CD11b^+^Ly6C^+^Ly6G^+^), dendritic cells (CD11b^+^Ly6C^-^Ly6G^-^CD24^+^), or macrophages (CD11b^+^Ly6G^-^Ly6C^-^F4/80^+^CD64^+^) (**Figure S3B-E**). We also found no common, significant changes in M1 (MHCII^high^CD206^low^) or M2 (MHCII^low^CD206^high^) macrophages or the ratio of M1/M2 (**Figure S3B-C**). The only difference observed in BPD6 model was a minor reduction in total macrophages and M1 macrophages (**Figure S3B**). In MC38, MR reduced total and MHCII presenting dendritic cells (**Figure S3E**). Regardless, MR shows no consistent impact on intratumor myeloid cells in four tested models.

We also evaluated lymphoid cell populations by probing for CD3, CD4, FoxP3 (marker for Tregs), CD8, CD44, CD69, IFNγ, and Granzyme B markers (**Figure S4A**). In BPD6 tumors, MR elevated the cell population expressing CD8^+^ T cell activation (CD44^+^CD69^+^) and cytotoxicity (IFNγ^+^) markers despite a slight reduction in the overall CD3^+^ T cells (**Figure 2A**). In YUMM5.2 tumors, MR led to a significant increase in total CD8^+^ T cells and activated CD8^+^ T cell population (IFNγ^+^ and Granzyme B^+^) (**Figure S4B**). Consistent with the syngeneic melanoma tumors, MR also increased the CD44^+^CD69^+^, IFNγ^+^, and Granzyme B^+^ CD8^+^ T cell populations in iBP tumors (**Figure S4C**). In line with the activated intratumor CD8^+^ T cells in the above melanoma models, MR resulted in an elevation in IFNγ^+^ and Granzyme B^+^ CD8^+^ T cell populations in MC38 tumors (**Figure 2B**). In addition, an increase in the repressive and exhaustive marker programmed cell death protein 1 (PD1) was found in YUMM5.2 model but not in the other three models. Furthermore, in all these models, MR showed no significant impact on CD4^+^ or regulatory T cells (Treg, CD4^+^FoxP3^+^) (**Figures 2A-B and S4B-C**). Overall, MR increases tumor infiltration of activated CD8^+^ T cells across different tumor models.

**Fig 2.**
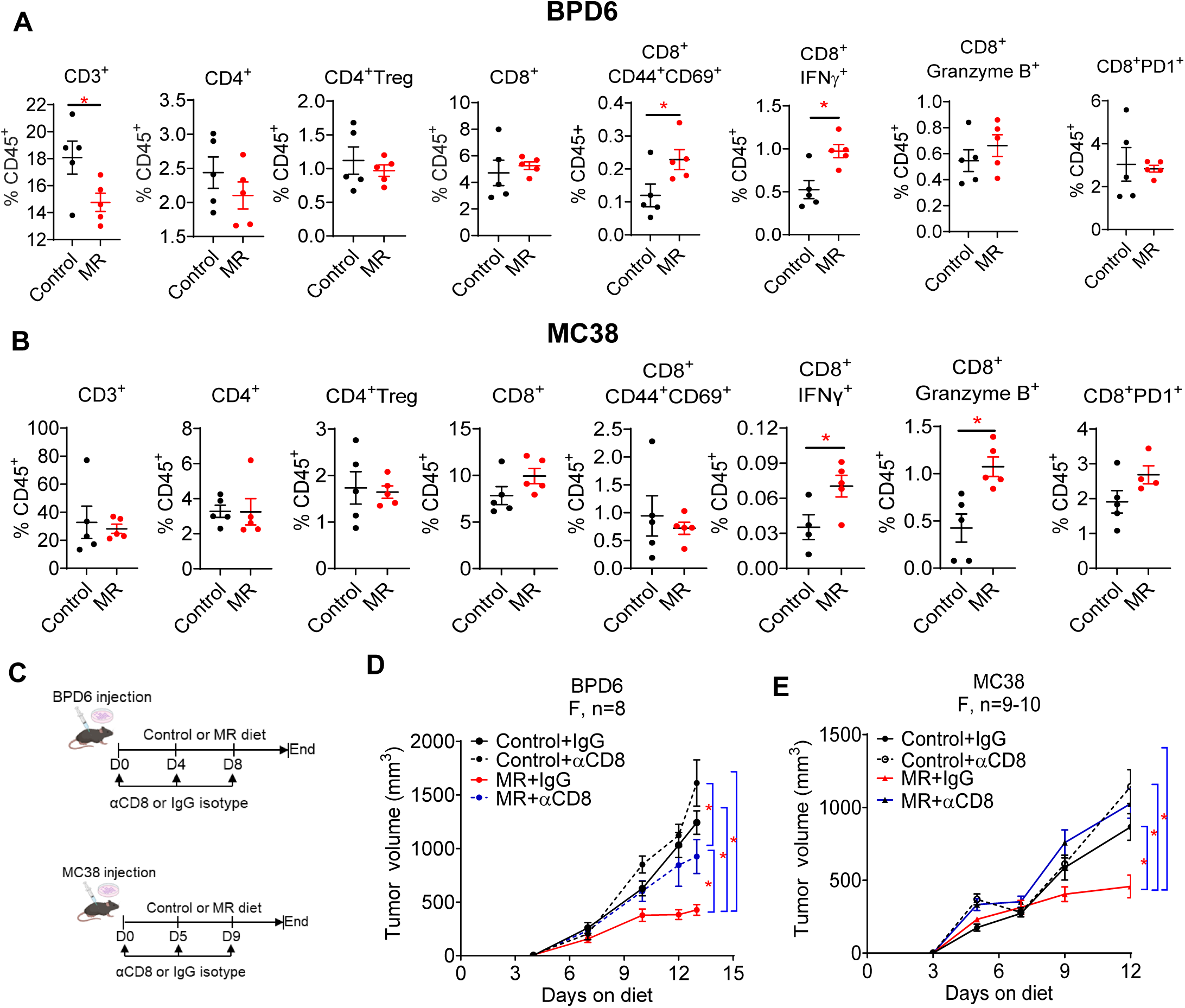
MR-inhibited tumor growth is CD8^+^ T cell dependent A. Flow cytometry analysis of lymphocytes in BPD6 tumors from female C57BL/6 mice. B. Flow cytometry analysis of lymphocytes in MC38 tumors from female C57BL/6 mice. C. Schematic for combination of dietary treatment, control or MR, with αCD8 or IgG isotype treatment in BPD6 and MC38 model. D. Tumor growth in BPD6 tumors. (*n =* 8/group) E. Tumor growth in MC38 tumors. (*n =* 9-10/group). Data were expressed as mean ± SEM. *P *<* 0.05, by two-tailed Student’s *t* test (A and B), and by 2-way ANOVA followed by Bonferroni’s multiple-correction test (E, F).

### MR-inhibited tumor growth is CD8^+^ T cell dependent

Since there was a consistent and significant elevation in intratumoral activated CD8^+^ T cells, we questioned whether MR-triggered tumor inhibition is CD8^+^-dependent. To address this, we inoculated mice with MC38 or BPD6 cells and then fed mice with the control or MR diet along with the treatment of αCD8 antibody or IgG isotype control every 4-5 days (**Figure 2C**). αCD8 antibody depleted intratumor CD8^+^ T cells in both BPD6 and MC38 models with no impact on CD3 or CD4 population (**Figure S4D-E**). CD8 depletion partially reversed MR-induced tumor inhibition in the BPD6 model and completely abolished MR-induced tumor inhibition in the MC38 model (**Figure 2D-E**). Furthermore, the anti-tumor effect of MR was abolished in MC38 tumors established in Rag1KO mice that lack functional T cells (**Figure S4F**). These data support that the anti-tumor effect of MR is CD8^+^ T cell-dependent, at least in BPD6 and MC38 models.

### Methionine is not limited in the tumor microenvironment (TME)

Nutrients in the TME are considered limited due to the high demand for malignant tumor growth^37,38^. Prior work in B16F10 melanoma model^28,29^ have shown that cancer cells have much higher expression of methionine transporter than T cells, which enables more efficient methionine uptake for tumor growth. Meanwhile, T cell activation is accompanied by significant uptake and metabolism of methionine^14,26,27^. Therefore, methionine was anticipated to be limited in the TME for the cytotoxicity of T cells. To determine whether methionine in the TME is limited in our experimental settings, we conducted metabolomics analysis on the tumor interstitial fluid (TIF) and the plasma from the same MC38 and YUMM5.2 tumor-bearing mice fed the control or MR diet (**Figure 3**). In both models, the metabolic profile in the TIF was distinct from that in the plasma, with some metabolites relatively higher in the TIF but others relatively lower when compared to the plasma (**Figure 3A-B and Figure S5A-B**). Consistent with previous analyses in pancreatic ductal adenocarcinoma and lung tumors^37^, glucose in the TIF was depleted in both MC38 and YUMM5.2 tumors regardless of the dietary treatment, which was accompanied with highly elevated lactate (**Figure 3A-C**), reflecting the highly glycolytic feature (Warburg effect) in both tumors. Interestingly in MC38, except for glutamine that was lower in the TIF (**Figure 3A and 3D**), levels of most other amino acids were either comparable (His, Try, and Arg) or higher (the remaining amino acids) in the TIF from control diet-fed mice than that in the plasma. In contrast, glutamate was ∼40-fold higher than plasma in MC38 (**Figure 3D**), indicating high glutamine consumption or high transamination in MC38 tumors. Similarly, in YUMM5.2 model, glutamate in TIF was ∼ 70-fold higher than in plasma, while glutamine levels remained the same (**Figure 3D**). Most of the other amino acids in the TIF from mice fed the control diet were also higher than that in the plasma or comparable, except for cystine, another sulfur-containing amino acid, which was 80% less than in the plasma (**Figure 3B and 3E**). Thus, unlike glucose and glutamine, methionine, along with most other amino acids, is not limited in the TIF compared to plasma.

**Fig 3.**
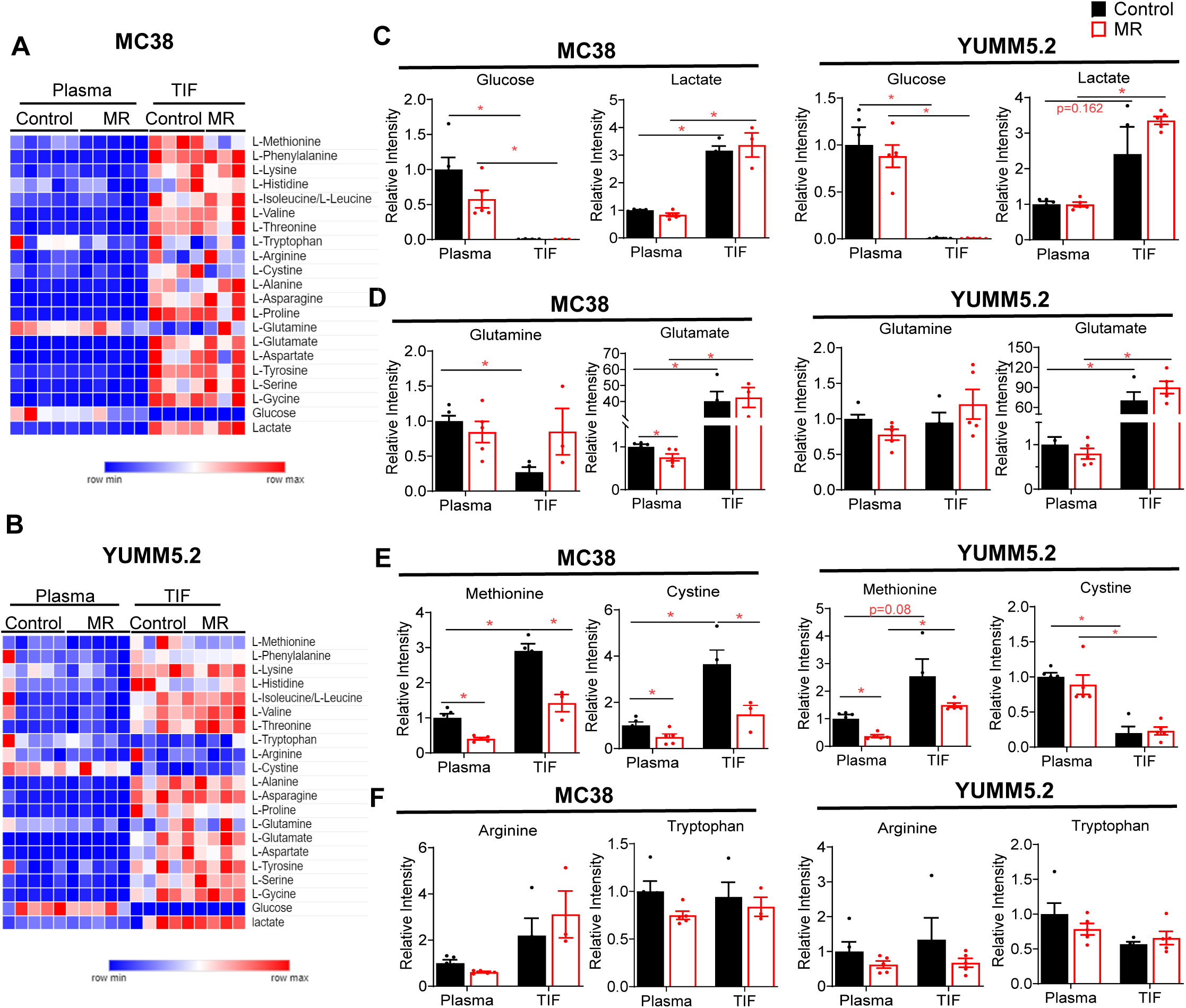
Methionine is not the limiting factor in the tumor microenvironment A. Heatmap of amino acids, glucose, and lactate in plasma and TIF from MC38 model. B. Heatmap of amino acids, glucose, and lactate in plasma and TIF from YUMM5.2 model. C. Relative mass intensity of glucose and lactate in plasma and TIF from MC38 and YUMM5.2 models. D. Relative mass intensity of glutamine and glutamate in plasma and TIF from MC38 and YUMM5.2 models. E. Relative mass intensity of methionine and cystine in plasma and TIF from MC38 model and YUMM5.2 models. F. Relative mass intensity of arginine and tryptophan in plasma and TIF from MC38 model and YUMM5.2 models. Data were expressed as mean ± SEM. *P *<* 0.05, by unpaired two-tail t-test. N *=* 5/group except for MC38-MR-TIF (*n =* 3) and MC38-Control-TIF (*n =* 4). TIF: Tumor interstitial fluid.

In MC38 model, MR reduced methionine and cystine in the plasma and TIF, and a slight reduction in Arg and Tyr in the plasma with no significant effect on other amino acids (**Figure S4A and Figure S5C-D**). Despite the MR-induced reduction, both methionine and cystine in the TIF remained comparable with that in the plasma from control-fed mice (**Figure 3E**). In YUMM5.2, MR decreased methionine levels in the plasma but not in the TIF and showed no effect on cystine in either plasma or TIF (**Figure 3E**). Similar to the MC38 model, methionine in the TIF remained comparable to that in the plasma from control-fed YUMM5.2 model (**Figure 3E**). The difference of metabolites in the plasma and TIF between the two models in response to MR may be attributed to the different genetics and cancer types since both models were established in the same sex and strain, female C57BL/6J mice. Furthermore, arginine and tryptophan, two other amino acids essential for T cell function^39^, were not changed by MR in the TIF from both MC38 and YUMM5.2 tumors (**Figure 3F**). Together, methionine, at least in the current contexts, is not limited in the TME.

### MR primes spleen CD8^+^ T cells for activation

Considering the significant increase in intratumor activated CD8^+^ T cells in various immune-competent models, we questioned whether MR alone affects the immune system *in vivo* in the absence of tumor. To this end, we first fed 7-8-week-old healthy C57BL/6J mice the control or MR diet as before for two weeks and then harvested the spleen and isolated splenocytes that were further immune stained for flow cytometry analysis (**Figure 4A**). We found that MR significantly increased CD3^+^ and CD8^+^ T cells, but not CD4^+^ T cells (**Figure 4B**), in the spleen of non-tumor-bearing mice. within CD3^+^ T cell population, CD4^+^ and CD8^+^ T cells are the two major cell types, both of which are important for tumor inhibition^40–42^. Since MR only increased CD8^+^, but not CD4^+^ T cells, we focused on CD8^+^ T cells and questioned whether such elevation is associated with alteration in cell metabolism, which is essential for T cell activation and function. To capture the metabolic features before and after activation, we performed metabolomics analysis on freshly isolated CD8^+^ T cells (basal condition) and activated CD8^+^ T cells (**Figure 4C**).

**Fig 4.**
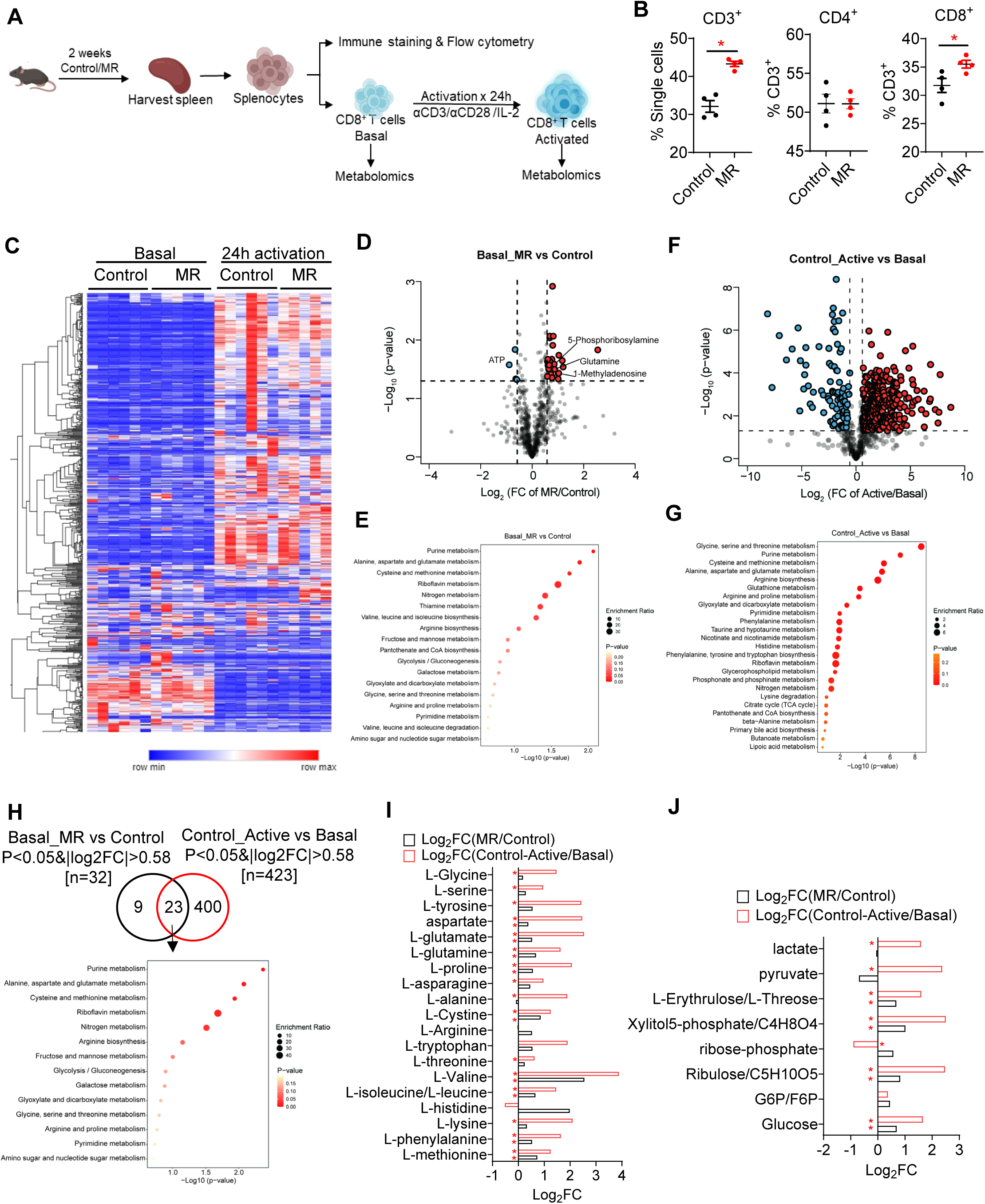
MR primes spleen CD8^+^ T cells for activation A. Workflow: C57BL/6J mice were fed the control or MR diet for two weeks, and splenocytes were isolated from the spleen and then stained for flow cytometry analysis or used for further isolation of CD8^+^ T cells that were either directly used for metabolomics analysis or cultured in the presence of αCD3CD28 antibody for 24 h before metabolomics analysis. B. Flow cytometry of single splenocytes isolated from mice fed the control or MR diet in A (*n =* 4/group). C. Heatmap of 684 metabolites in CD8^+^ T cells isolated from mice fed control or MR diet before and after activation (*n =* 5-6/group). D. Volcano plot of 684 metabolites in CD8^+^ T cells isolated from mice fed control or MR diet under basal condition before activation. Colored dot indicates that P < 0.05 & |log2FC| > 0.58. E. Gene Ontology analysis of significantly changed (*P* < 0.05 & |log2FC| > 0.58) genes by MR in D. F. Volcano plot of 684 metabolites in CD8^+^ T cells isolated from mice fed control diet before and after activation. Colored dots indicated metabolites with *P* < 0.05 & |log2FC| > 0.58. G. Gene Ontology analysis of significantly changed (*P* < 0.05 & |log2FC| > 0.58) genes by MR in F. H. Venn diagram of commonly changed (*P* < 0.05 & |log2FC| > 0.58) metabolites in D and F; and Gene Ontology analysis of the 23 commonly changed metabolites. I. log2FC of amino acids in Dand F. J. log2FC of metabolites in glycolytic pathway in Dand F. Data were expressed as individual data points and indicate the mean ± SEM. *P *<* 0.05, and ^0.05 < P *<* 0.1, by two-tailed Student’s *t* test (B).

Under the basal conditions, MR significantly altered (*P* < 0.05 & |log_2_FC|>0.58) 32 intracellular polar metabolites, and further pathway analysis revealed the top metabolic hits of MR on purine metabolism, followed by amino acid metabolism and riboflavin metabolism (**Figure 4D-E and Figure S6A**). In purine metabolism pathway, ATP was significantly reduced while L-glutamine, and 5-phosphoribosylamine were increased (**Figure 4D**), suggesting an upregulation of purine biosynthesis and a high energy demanding state. For T cell activation, CD8^+^ T cells were cultured in the presence of αCD3 and αCD28 antibodies and cytokine IL-2 for 24 h. T cells are known to reprogram their metabolism upon activation. As anticipated, activation led to a striking alteration of cellular metabolites in CD8^+^ T cells from both control and MR-fed mice with amino acid metabolism, and purine and pyrimidine metabolism being the most impacted pathways (**Figure 4F-G and Figure S6B-C**). After activation, the difference between cells from control-fed and MR-fed mice was minimal with only a few metabolites significantly changed (**Figure S6D-E**). Notably, 23 metabolites were commonly changed by MR under basal condition and by activation in control CD8^+^ T cells, and these metabolites were mapped to purine metabolism and amino acid metabolism, followed by riboflavin and glucose metabolism as the top altered metabolic pathways (**Figure 4H**). Indeed, under basal state, the majority of amino acids were either significantly increased or with a trend towards increase except alanine, and such increase was largely recapitulated and magnified by T cell activation (**Figure 4I**). The MR-induced changes in purine metabolism under basal condition were replicated by activation (**Figure S6A**). Similarly at basal condition, glucose and other intermediates in the glycolytic pathway, G6P/F6P (glucose 6-phosphate/fructose 6-phosphate), Ribulose/C5H10O5, xylitol5-phosphate/C4H8O4, and L-Erythrulose/L-Threose, were also significantly increased showing a trend towards increase by MR while cellular lactate, which was likely secreted out of cells, remained comparable (**Figure 4J**), suggesting an elevated glycolytic pathway, another feature of activated T cells^43^. Again, the changes in glycolytic pathways are largely magnified by activation (**Figure 4J**). The MR-induced increase in riboflavin, however, was reduced by activation (**Figure S6A**).

Interestingly, cellular methionine and cystine levels were significantly increased in CD8^+^ T cells from MR-fed mice, while other methionine-related metabolites, SAM, SAH, 5-methylthioadenosine (MTA), and SAM/SAH ratio, remained the same (**Figure S6F**), suggesting minimum impact on cellular methylation reactions. Overall, through metabolomics, MR appears to prime spleen CD8^+^ T cell toward activation through purine metabolism, amino acid metabolism, and glycolytic pathway.

Furthermore, we performed RNA sequencing (RNA-seq) analysis on CD8^+^ T cells under the basal condition. MR significantly (*Padj* < 0.05 & |log_2_FC|>1) up-regulated 108 genes, and down-regulated another 108 genes (**Figure S7A-C**). Gene Ontology (GO) analyses of 108 significantly up-regulated genes revealed that MR significantly enriched genes in the NOTCH3 signaling, which has been shown to be important for T cell lineage specification and regulatory T cell homeostasis and function^44,45^, and MHCII antigen presentation (**Figure S7B**), which is the cornerstone of T cell recognition and function^46^. In line with the metabolomics data, GSEA analysis showed that MR led to a significant increase in glucose and amino acid metabolism related pathways (**Figure S7D**). In glucose metabolism, MR resulted in a significant increase in *H6pd (Hexose-6-Phosphate Dehydrogenase), Pdk* (pyruvate dehydrogenase), *Eno1* (Enolase 1), *Ldhb* (lactate dehydrogenase b subunit), *Pygm* (glycogen phosphorylase), *Galk1* (galactokinase), and *Acss2* (acetyl-coenzyme A synthase) **(Figure S7E).** Notably, we and others have shown that *Eno1* encodes a glycolytic enzyme essential in sustaining the cytotoxicity effect of tumor infiltrated T cells^47,48^. In amino acid metabolism, MR significantly increased three aldehyde dehydrogenase family members (*Aldh6a1*, *Aldh1l2*, and *Aldh1A1*) that are involved in aldehyde metabolism and one carbon metabolism, Glutathione S-Transferases (*Gstt1* and *Gstp2*), and *Nfe2l2* (Nuclear Factor Erythroid 2–Related Factor 2) that regulates cellular redox balance (**Figure S7F**). On the other hand, MR also significantly downregulated *Gad1* (glutamate decarboxylase 1), *Hal* (Histidine Ammonia-Lyase), and *Hdc* (histidine decarboxylase), all of which are related to amino acid catabolism (**Figure S7F**). Together, RNA-seq data further support the metabolic impact of MR on glucose and amino acid metabolism, in addition to other potential mechanisms through signaling and antigen presenting.

### MR enhances the efficacy of αPD1 immunotherapy

We further evaluated the effect of MR in combination with immune checkpoint inhibitor αPD1 immunotherapy (**Figure 5A-B**). αPD1 effectively reduced PD1^+^CD8^+^ T cells in BPD6 model, and to a less extent in MC38 model (**Figure 5C-D**). The less reductive effect on the PD1^+^CD8^+^ T cells in MC38 model could be due to the relatively higher population of CD8^+^ T cells overall in MC38 (7.83%) than in BPD6 (4.71%) (**Figure 2A-B**). αPD1 alone slowed down tumor growth in MC38 model but not in BPD6 model. With the treatment of control IgG isotype antibody, MR-mediated tumor inhibition remained in MC38 model but not in BPD6 model (**Figure 6E-F and Figure S8A-B**). The lack of tumor inhibition by MR in the BPD6 model here could be attributed to the IgG treatment per se, which is known to affect the immune system in both spleen and peripheral tissue^49^. Importantly, MR and αPD1 resulted in a most striking tumor inhibition in both models (**Figure 6E-F and Figure S8A-B**). Together, MR improves the efficacy of anti-PD1 immunotherapy in BPD6 and MC38 models.

**Fig 5.**
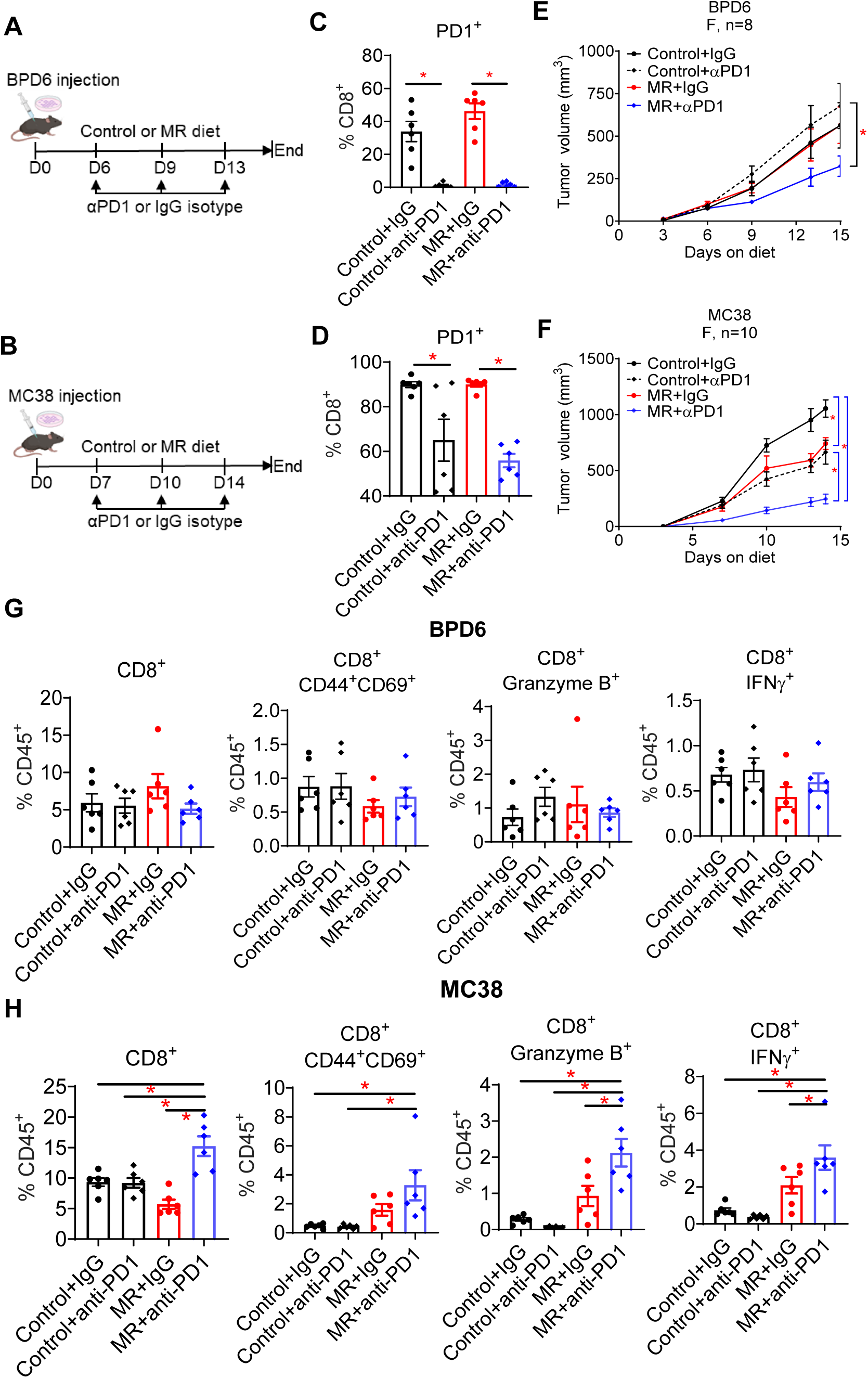
MR enhances efficacy of anti-PD1 immunotherapy A. Workflow for BPD6 model treated with control or MR diet in combination with IgG isotype control or anti-mouse PD1 (αPD1) antibody (100 µg/mouse/dose). B. Workflow for MC38 model treated with control or MR diet in combination with IgG isotype control or αPD1 antibody (100 µg/mouse/dose). C. BPD6 intratumor PD1^+^CD8^+^ T cells analyzed by flow cytometry (*n =* 5/group). D. MC38 intratumor PD1^+^CD8^+^ T cells analyzed by flow cytometry (*n =* 5/group). E. Tumor growth in BPD6 model (*n =* 8/group). F. Tumor growth in MC38 model (*n =* 10/group). G. Flow cytometry analysis of intratumor T cell markers in BPD6 tumors. (*n =* 5/group). H. Flow cytometry analysis of intratumor T cell markers in MC38 tumors. (*n =* 5/group). Data were expressed as mean ± SEM. *P < 0.05 by 2-way ANOVA followed by Bonferroni’s multiple-correction test for C-F, and by one-way ANOVA test for G&H.

To determine whether the enhanced anti-tumor effect was due to activated intratumor CD8^+^ T cells, we collected BPD6 and MC38 tumors at the endpoint for flow cytometry analysis. αPD1 alone showed no significant impact on the CD3^+^, CD4^+^, or Treg populations in either BPD6 or MC38 model (**Figure S7C-D**). αPD1 alone also had no significant effect on CD8^+^ T cell activation in both BPD6 and MC38 models (**Figure 5G-H**). MR-induced intratumor CD8^+^ T cell activation was abolished in BPD6 model with IgG control treatment (**Figure 5G**). The combination of MR and αPD1 also led to no significant change in CD3^+^, CD4^+^, Treg^+^, CD8^+^T cells or activated CD8^+^ T cells (CD44^+^CD69^+^, Granzyme B^+^, and IFNγ^+^) in BPD6 tumors (**Figure 5G and Figure S7C**), suggesting factors other than CD8^+^ T cells may partake in the tumor inhibition observed in this model. On the contrary, in MC38 tumors, the combination of MR and αPD1 significantly increased CD3^+^, and CD8^+^ T cell populations as well as the activated T cell markers, CD44^+^CD69^+^, Granzyme B^+^, and IFNγ^+^ (**Figure 5H and Figure S7D**). Thus, in MC38 model, the additive effect of MR and αPD1 is attributed to increased CD8^+^ T cell population and activation. Together, the mechanisms underlying the MR-improved efficacy of αPD1 immunotherapy appear to be complex and context-dependent.

## Discussion

We provide compelling evidence supporting the anti-tumor effect of MR in an immune-competent mouse background through seven syngeneic mouse models for melanoma, colorectal cancer, breast cancer, and one genetically modified melanoma mouse model. The syngeneic mouse models were established in 3 different mouse stains, C57BL/6J (for BPD6, YUMM5.2, MC38, A7C11 and LLC1 models), and FVB (for Met-1 model). They also carry distinct genetic features: BPD6^50^ and iBP^32^ are *Braf^V600E^Pten^−/−^*, YUMM5.2^51^ carrying *Braf^V600E^p53^−/−^ Cdkn2^+/-^*, MC38 carrying Trp53 mutations (G242 V and S258I)^52^, A7C11^53^ carrying *p53^−/−^/Kras^G12D/+^*, Met-1 being a triple negative breast cancer line derived from a FVB/N-Tg(MMTV-*PyVmT*) tumor^54^, and hypermutated LLC1 with mutations present in 30 genes including *Kras*, *Nras* and *p53*^55^. MR alone led to a significant tumor inhibition in BPD6 (female), YUMM5.2 (female and male), iBP, MC38 (male), A7C11, Met-1, and LLC1 (female) models, and a trend toward inhibition in BPD6 (male) and MC38 (female), where statistical significance would likely be reached with a larger sample size. Indeed, recent studies from others^20,21,30^ have also shown the anti-tumor effect of MR in MC38 and CT26 models established either subcutaneously or orthotopically in an immunocompetent background. Considering the wide range of mouse genetic backgrounds and cancer cell genetic mutations, the anti-tumor effect of MR does not appear to be mouse strain-dependent or solely RAS mutation-dependent^7^. Furthermore, the anti-tumor effect is not sex-dependent since the tumor inhibition effect was present in males and females in 3 different models, BPD6, YUMM5.2, and MC38.

Our data is consistent with a few recent reports supporting the anti-tumor effect of MR in immunocompetent mice. The anti-tumor effect of MR in melanoma has also been recapitulated in B16F10 melanoma model^21^. In the same model, a tumor-suppressing effect can be achieved via alternative methionine-lowering approaches, either by intermittent methionine deprivation^56^ or by intravenous injection of a genetically modified bacterial strain *Salmonella typhimurium* that overexpresses an L-methioninase and prefers targeting tumor for methionine reduction^57^. For colorectal cancer, a significant MR-induced tumor inhibition has been reported in both MC38^19–21^ and CT26^20^ models that are established via subcutaneous injection. Similar tumor inhibition has also been reached in MC38 and CT26 models established via cecal orthotopic tumor inoculation^30^. Here, we provide new evidence supporting an anti-tumor effect of MR in two immune-competent breast cancer models, A7C11 and Met-1. Moreover, a beneficial anti-tumor effect of MR has also been shown in immunocompetent non-small lung cancer mouse model driven by *Kras^G12D^Lkb1^-/-^*^22^, glioblastoma^23^, *Myc*-driven prostate, and renal cell carcinoma models^24^. Our current report, adding to accumulating evidence, supports that MR via dietary intervention can impede tumor growth in a wide range of immune-competent, preclinical (syngeneic and genetic) mouse models.

Does the immune system contribute to MR-induced tumor inhibition? We have addressed this question by first evaluating dietary MR effect in immune-deficient NSG mice bearing BPD6, YUMM5.2, MC38, or Met-1 tumors. Tumor inhibition was completely abolished in YUMM5.2 and MC38 tumors and appeared to be partially reversed for BPD6 and Met-1 tumors in NSG mice, suggesting the anti-tumor effect is attributed to the immune system, and the extent of dependency is context-dependent. Further flow cytometry analysis in BPD6, YUMM5.2, MC38, and iBP models revealed that one common change is the activation of intratumor CD8^+^ T cells reflected by elevated activation marker CD44^+^CD69^+^, and cytotoxicity markers IFNγ^+^ and Granzyme B^+^ (**Figure 2 and S4**). No common significant impact was found in PD1^+^CD8^+^ T cells, which was elevated in YUMM5.2 tumors, but not in BPD6, MC38, or iBP (**Figure 2A-B and Figure 2B-C**), CD4^+^ T cells, Treg, DCs, monocytes, neutrophils, or macrophages (M1 and M2) (**Figure S3**). The absence of tumor inhibition in MC38 tumors established in Rag1KO mice (Figure S2F) consolidates that MR-inhibited tumor growth in this model is primarily attributed to CD8^+^ T cells. In line with our findings, an increased CD8^+^ T cell population or activation has been found by others in colorectal cancer models MC38 and CT26^20,21,30^, B16F10 melanoma model^21^, and in RP-B6-Myc prostate model^24^. MR caused by intermittent methionine deprivation also increases CD8^+^ T cell population in B16F10 tumors^56^. Considering the crucial role of CD8^+^ T cells in immune defense, tumor surveillance, and killing^41,58^, it’s not surprising that MR improves the efficacy of immune checkpoint blockades αPD1 (**Figure 5**), αPD-L1(programmed death-ligand 1), and αCTLA-4^19–21,24,30,56^. Thus, CD8^+^ T cells appear to be the key to MR-induced tumor inhibition and MR is a potential booster for immune checkpoint blockades.

How does MR increase intratumor CD8^+^ T cells and boost the response to immune therapy? Prior studies have discovered multiple tumor cell-intrinsic mechanisms. For instance, MR triggers a dramatic reduction in cellular SAM level that decreases histone methylation marks in tumor cells and tumor-initiating cells that reduce tumor growth and tumorgenecity^8,18^. Li et al. has shown in tumor cells that MR led to a reduction in N6-methyladenosine (m6A) methylation, not histone methylation or 5-methylcytosine DNA methylation, thereby downregulating the translation of immune checkpoints PD-L1 and V-domain Ig suppressor of T cell activation (VISTA)^20^. MR also decreases SAM-mediated methylation of cyclic GMP-AMP synthase (cGAS) in tumor cells, which further activating the cGAS-STING signaling pathway for IFN production^19,21^. The anti-tumor effect of MR has also been linked to its contribution to redox balance. MR sensitizes tumor cells to ferroptosis due to its contribution to GSH synthesis through the transsulfuration pathway and GSH degradation^23,56^. Interestingly, intermittent methionine deprivation enhances ferroptosis via upregulation of endoplasmic reticulum (ER) stress through the eIF2α/ATF4 pathway, initiating the transcription of downstream CHAC1 and CHAC1-mediated GSH degradation, with no impact on protein synthesis, whereas prolonged methionine deprivation desensitizes response to ferroptosis through downregulation of CHAC1^56^. Similarly, general protein restriction also increases immune surveillance through a tumor-specific IRE1a meditated ER stress, but not in tumor-infiltrated lymphocytes or DCs^59^. Together, existing evidence suggests that the increased intratumor CD8^+^ T cells may be attributed to its tumor intrinsic effect on methylation reaction, GSH synthesis, and ER stress.

In addition to the tumor autonomous mechanism, nutrient availability and cell-cell interaction in the TME are critical in tumor progression and treatment^60,61^. Methionine-centered one-carbon metabolism is indispensable for CD4^+^ and CD8^+^ T cell activation and function^14,25,26^. Since both tumor cells and T cells require methionine for their growth and function, the anti-tumor immunity then depends on the availability of methionine in the TME and the capability of uptake. Recent studies suggested tumor cells in the TME could outcompete T cells for methionine uptake via transporter SLC43A2, resulting in T cell exhaustion via SAM-mediated H3K79me2 histone methylation^28,29^. On the other hand, tumor methionine metabolism also drives T cell exhaustion in hepatocellular carcinoma via tumor-derived SAM and MTA that reprograms CD8^+^ T cell metabolism, leading to T cell dysfunction^62^. Interestingly, by probing for metabolite availability in the TME (TIF), we found that the majority of amino acids, including methionine, in the TIF are not low compared to plasma. Although MR decreased (significantly or trending) methionine in the circulation and TIF, levels in the TIF remained higher than that in the plasma from control-fed mice (**Figure 3**), suggesting that methionine is not the limiting factor in the TME, at least in the tested models. This is unexpected considering previous studies have suggested methionine deficiency in the TME of the B16F10 melanoma model ^28,29^. This discrepancy may be due to different melanoma tumor models and their distinct genetic backgrounds. Unlike the melanoma models used here, the prevalent BRAF mutation in human melanoma is not present in the B16F10 model^52^, despite being widely used. Considering the partitioning of nutrients in the TME may be defined by cell-intrinsic mechanisms^63^, future studies are warranted to define the uptake and metabolism of methionine by the tumor and different immune cells in the TME.

Another intriguing finding is that MR increases the population of CD3^+^ and CD8^+^ T cells from spleen of non-tumor bearing mice, and primes CD8^+^ T cell metabolism for activation. In the metabolite level, purine metabolism and amino acid metabolism are the top hits by MR, followed by glycolysis pathway. The changes in all these three pathways are replicated or magnified by T cell activation, implying a primed activation status. In addition, other mechanisms such as NOTCH3 signaling, and MHCII antigen presentation may also contribute to T cell priming and activation that await future investigation. Indeed, methionine-derived H_2_S inhibits CD8^+^ T cell migration^30^. With these data, we speculate that MR primed immune surveillance at the systemic level to better prepared to inhibit tumor progression.

It is noteworthy to mention that our current study is different from another recent study by Ji et al.^31^, which is very different from ours and others^19–21,30^ despite the same models and the same or similar diets used. The difference cannot be explained by the location of tumor as both orthotopic and subcutaneous injections were explored by others. One possible explanation is the difference in the initial point for dietary treatment and the duration of the treatment. Unlike all the other studies where dietary intervention was initiated post-tumor injection or concurrently, Ji et al. started dietary treatment 3-4 weeks prior to tumor injection^31^, resulting in a longer overall treatment duration compared to these other studies, including ours. The duration of MR and the extent of methionine restriction could determine the therapeutic effect through distinct mechanisms^56^.

In summary, MR impedes tumor growth in a wide range of immune-competent, preclinical mouse models and is a potential booster for immune checkpoint blockades. MR-induced tumor inhibition may depend on CD8^+^ T cells in a context-dependent matter. Mechanistically, MR alters spleen T cell populations in the systemic level and primes T cell metabolism for activation. Together, our data provides a rationale for employing dietary MR as an adjuvant therapy along with immune checkpoint inhibitors and other therapeutics to promote anti-tumor immunity.

## Method

### Animals and diets

All animal procedures and studies were approved by the Institutional Animal Care and Use Committee (IACUC) at Duke University (Durham, North Carolina, USA) or Baylor College of Medicine (Houston, Texas, USA). All experiments were performed in accordance with relevant guidelines and regulations. All mice were housed at 20 ± 2°C with 50 ± 10% relative humidity and a standard 12 h dark-12 h light cycle. The special amino acid-defined diets (Supplementary Table I) were designed with minor modifications from the diets used before^7^ with the control diet containing 0.86% methionine (w/w, catalog #510027) and the MR diet containing 0.17% methionine (w/w, catalog #511405). This minor increase of methionine in the MR diet is based on the recent report showing that 0.17% of methionine led to no impact on body weight gain or lean mass, but reduced fat mass^33^. All amino acid-defined diets were made by Dyets Inc. (Bethlehem, PA, USA) and were irradiated before use. C57BL/6J (JAX:000664), FVB (JAX: 001800), BALB/c (JAX:000651), NSG (NOD.Cg-*Prkdc^Scid^ Il2rgtm1Wjl*/SzJ, JAX:005557) and Rag1KO (JAX:002216) mice were purchased from the Jackson Laboratory or the Center for Comparative Medicine at Baylor College of Medicine. Age-matched mice at 7-8-week-old were used for all the studies. Mice had ad libitum access to food and water.

We evaluated MR effects on healthy (non-tumor-bearing) mice and a variety of tumor bearing mice. For all dietary and tumor experiments, mouse body weight and tumor size were monitored 2-3 times per week until the end of the study. Mice were euthanized by CO_2_ when there was more than 20% body weight loss compared to their initial body weight, or the tumor size exceeded 1.5 cm^3^.

### Non-tumor bearing mice

7-8-week-old male C57BL/6J mice were randomized to either the control or MR diet ad libitum for 2 weeks. Mouse spleens were collected for isolation of splenocytes, CD3^+^ T cells, and CD8^+^ T cells detailed below.

### Syngeneic mouse models

For syngeneic melanoma models, BPD6 (5 x 10^5^) and YUMM5.2 (5 x 10^5^) cells were injected subcutaneously into C57BL/6J mice for tumor induction. For syngeneic colorectal cancer models, MC38 (5 x 10^5^) cells were injected into the flank of C57BL/6J mice. For syngeneic breast cancer models, A7C11 (2 x 10^5^) and Met-1 (2 x 10^5^) cells were injected subcutaneously into the third mammary fat pad of C57BL/6J mice and FVB mice, respectively. For lung cancer models, LLC1 (5 x 10^5^) and Met-1 (2 x 10^5^) cells were injected subcutaneously into the third mammary fat pad of C57BL/6J mice and FVB mice, respectively. For all syngeneic mouse models, 7-8-week-old mice were used for tumor cell inoculation and subjected to the control or MR diet on the same day, right after cell injection, continuing until the endpoint. Both females and males were used for BPD6 and YUMM5.2 models, and female mice were used for the other models. Tumors were measured thrice weekly using an electronic caliper. Tumor volumes were measured by the formula V=L x W^2^/2, where L is length and W is width.

### Tumor growth in immune-compromised NSG and Rag1KO mice

To evaluate the effect of MR in immune-compromised background, female NSG mice were injected with BPD6, YUMM5.2, MC38, and Met-1 cells as outlined in the immune-competent mice. Mice were randomized to the control or MR diet immediately after cell injection. To confirm the relevant of T cells in MR-mediated tumor inhibition, female Rag1KO mice were injected with MC38 cells as before and then randomized to the control or MR diet immediately after cell injection.

### Genetically modified mouse model

iBP (*BrafV600E/WT*, *Ptenfl/fl* mTyrCreERT2) mice were generated by crossing *BrafWT/WT Ptenfl/fl* mTyrCreERT2 mice with *Braf^V600E^ Pten^fl/fl^* mice. Tumor was induced with a single intradermal dose of 4-hydroxytamoxifen in DMSO (150 µg/mouse). The latency for tumor development in these animals was typically ∼ 21-28 days. Tumors were measured thrice weekly using an electronic caliper. Tumor volumes were measured by the formula V=L x B x H. The mouse melanoma cell line BPD6 was established from iBP as described^32^.

### αPD1 immunotherapy treatment

Age-matched C57BL/6J mice harboring MC38 tumor or BPD6 tumor were treated with αPD1 (clone: 29F.1A12, BioXCell, Lebanon, NH) or rat IgG2a (clone 2A3, BioXCell, Lebanon NH) at 100 µg/mouse (diluted in sterile PBS) by intraperitoneal injections starting at day 6 or day 7 post tumor cell injection and every 3 or 4 days after until the endpoint is reached.

### T cell depletion with αCD8 antibody

C57BL/6J mice were inoculated with MC38 or BPD6 cell, then subjected to the control or MR diet immediately after cell injection. For CD8^+^ depletion, mice were injected intraperitoneally with 200 µg/mouse of rat anti-CD8a antibody (clone YTS169.4, BioXCell) or rat IgG2b-anti-KLH isotype control (clone LTF2, BioXCell) diluted in sterile PBS, right before tumor cell injection and every 4-5 days after cell injection. The efficiency of CD8a depletion was analyzed at the end of the experiment by flow cytometry analysis of T cell subpopulations on blood obtained via cardiac puncture.

### Cell line and cell culture

The mouse cell lines YUMM5.2, and LLC1 were purchased from American Type Culture Collection (ATCC). BPD6 cells were a gift from Dr. Brent A. Hanks (Duke University). MC38 cells were a gift from Dr. Jatin Roper. A7C11 cells were provided by Dr. J. Conejo-Garcia (Duke University). Met-1 cells were obtained from Dr. Alexander Borowsky (University of California at Davis) and grown in Dulbecco’s Modified Eagle Medium (DMEM) with the same supplements. All cell lines were routinely tested for mycoplasma. All cells were cultured at 37 °C with 5% CO_2_ in RPMI 1640 media with the addition of 10% fetal calf serum, 100,000 U/L penicillin, and 100 mg/L streptomycin. Cell lines were authenticated at the Duke University DNA Analysis Facility by analyzing DNA samples from each cell line for polymorphic short tandem repeat (STR) markers using the GenePrint 10 kit from Promega (Madison, WI, USA). All cell lines were negative for mycoplasma contamination.

### Single-cell isolation from tumors

Tumors from syngeneic models were isolated, minced on a petri dish with media (DMEM + 5% FBS) and then enzymatically digested for 30-45 min with the addition of 100 µg/ml DNase I (D5025-150KU, Sigma-Aldrich, Denver CO) and 1 mg/ml collagenase (Collagenase A, Cat# 10103586001, Sigma-Aldrich, Denver CO). For iBP model, tumors were sliced into large chunks and subjected to mechanical digestion in a GentleMACS dissociator for 30 seconds. Following this, tumors were digested with an enzyme cocktail containing the above DNase I, collagenase, and 100 µg/ml hyaluronidase (H6254, Sigma-Aldrich, Denver CO) for 40 min, followed by a second round of mechanical digestion for 30 seconds. The cells were then filtered with a 40 µm strainer to produce single-cell suspensions, and the enzymes were diluted by the addition of media. Red blood cells were lysed with the addition of ACK lysis buffer (Cat #A1049201, Thermo Fisher Scientific, Waltham, MA) for 4 min at room temperature. Following red blood cell lysis, cells were washed with PBS before proceeding to flow cytometry or magnetic bead-based isolation.

### Single cell isolation from spleen

Spleens were collected from non-fasted mice after a 2-week feeding on either control or MR diet. Each spleen was prepped through gentle mashing with a syringe plunger in 5 ml of PBS containing 2% FBS and 1 mM EDTA and filtered through a 40 μm cell strainer (Falcon, 352340). The cell suspension was centrifuged (1000 g × 4 mins at 4°C) and supernatant was discarded. Cells were then resuspended in 2 ml ACK lysing buffer (Life Technologies, A1049201) for 3 min at 4°C to remove red blood cells. The lysis was stopped by addition of 6 ml PBS+2% FBS+1mM EDTA followed by centrifugation. Cell pellets were resuspended in 1 ml PBS containing 2% FBS and 1 mM EDTA for cell counting, and cells were redistributed for T cell isolation and flow cytometry staining and analysis.

### CD8^+^ T cell isolation and culture

CD8^+^ T cells were isolated from the splenocytes obtained above from mice fed the control or MR diet using the EasySep™ Mouse CD8^+^ T Cell Isolation Kit (Stem Cell Technologies, 19853) following manufacturer’s protocol. 1 million freshly isolated CD8^+^ T cells were pelleted and snap frozen as the basal condition for in-house metabolomics analysis or RNAseq analysis (Novogene Sequencing Services). CD8^+^ T cells were cultured in anti-CD3 (0.5 µg/ml)-coated 96-well plates in RPMI medium containing 8% fetal bovine serum, 1% non-essential amino acids, 1% 100 mM sodium pyruvate, 1% 1 M HEPES, 0.1% 50 mM 2-mercaptoethanol, 50 U/mL penicillin, and 50 µg/mL streptomycin, and activated by addition of 1.0 µg/mL anti-CD28 antibody and 10ng/mL IL-2. 24 h post activation, 0.4 million active CD8^+^ T cells were collected for metabolomics analysis.

### Flow cytometry staining

Single-cell suspensions (10^6^ cells in 100ul) from tumors were incubated with Live/dead fixable dead cell stain in PBS for 10 min at 4°C. Cells were spun down at 1000 g × 5 min and were incubated with Fc blocker anti-CD16/32 (Catalog# 14-0161-85, Thermo Fisher Scientific, Waltham MA) in flow buffer (10gms BSA in 1 L PBS, pH7.2) for 15 mins. Following this, cells were stained with an antibody cocktail in BV buffer (Cat# 566349, Thermo Fisher Scientific). The antibodies used are listed in Supplementary Table II. For intracellular staining, cells were fixed and permeabilized using transcription factor staining kit buffer (Cat# 00-5523-00, Thermo Fisher Scientific, Waltham, MA) followed by intracellular staining with desired antibody for 30 minutes at 4°C. CD3, CD4, FoxP3, CD8, CD44, CD69, IFNγ and Granzyme B markers, and myeloid cell population via probing against CD11b, CD11c, CD24, CD64, F4/80, MHCII, and CD206 markers. Cells were washed twice in Perm/Wash buffer and then placed in staining buffer for flow cytometry analysis.

Splenocytes (2×10^6^ per well) from non-tumor-bearing mice fed the control or MR diet were first stained with a fixable viability dye (Zombie UV, Biolegend). Cell surface markers were then stained using the following antibodies: anti-CD3-FITC (Clone 145-2C1, Biolegend), anti-CD8-BV605 (Clone 53-6.7, Biolegend), anti-CD4-PE Dazzle (Clone RM4-5, Biolegend). Following surface marker staining, cells were fixed and permeabilized using eBioscience™ Foxp3/Transcription Factor Staining Buffer Set (Invitrogen), then stained with anti-IFNγ-APC (Clone XMG1.2, Biolegend) and anti-Granzyme B-PE (Clone QA16A02, Biolegend).

For validating the efficiency of αCD8 treatment, blood collected via cardiac puncture at the endpoint was collected for staining for CD3, CD4, and CD8 markers as described above for the cells derived from tumors.

Multicolor flow cytometry was performed in BD Fortessa 16 color analyzer. Gating cell populations was done using isotype and single stain controls. Gating strategies and which figures they correspond to are outlined in Figure S2 and S3. The FACS results were analyzed by FlowJo_V10 software (FlowJo, LLC).

### RNA Sequencing

Total RNA was isolated from freshly isolated CD8^+^ T cells using QIAzol (QIAGEN, Germantown, MD, USA). The library construction, following a non-directional poly-A tail selection, and the following RNA sequencing was conducted by Novogene. Libraries were sequenced using a NovaSeq 6000 System (Illumina) using 20 million paired-end 150 bp (PE150) reads sequencing. Gene-set enrichment analysis (GSEA) on RNA-seq data was conducted using the gage function and non-parametric Kolmogorov–Smirnov test from the GAGE (version 2.22.0) R Bioconductor package.

### Metabolite Extraction

The extraction of cellular metabolite was similar as before (Gao et al., 2018, Cell Reports 22, 3507–3520). 4 × 10^5^ CD8^+^ T cells were counted, pelleted by centrifugation, and then snap frozen for metabolite extraction. 1 mL of ice-cold extraction solvent (80% methanol/water) and silica beads were added to each sample and samples were homogenized using a TissueLyser II (Qiagen). The extractant was centrifuged at 20,000 x g for 10 min at 4°C. 400 µL of supernatant was transferred to a new Eppendorf tube and dried in a vacuum concentrator. For spent media, 10 μl was used for extraction using 1 mL ice-cold 80% methanol/water via vortexing at the medium setting on a benchtop vertex for 1 min, and 400 µL of supernatant was dried for further analysis. For plasma and TIF, 1 μl was used for extraction using 500 µL ice-cold 80% methanol/water via vertexing for 1 min, and 200 µL of supernatant was dried in a vacuum concentrator. The dry pellets were stored at-80°C for LC-MS analysis. Samples were reconstituted into 30 µL of sample solvent (water:methanol:acetonitrile, 2:1:1, v/v/v) and were centrifuged at 20,000 x g at 4°C for 3 min. The supernatant was transferred to LC vials for analysis, and the injection volume was set at 3 μl.

### High-Performance Liquid Chromatography-Mass Spectrometry

Chromatography separations were carried out using a hydrophilic interaction chromatography method (HILIC) with an Xbridge amide column (100 × 2.1 mm i.d., 3.5 μm; Waters) on the Vanquish Horizon UHPLC system (Thermo Scientific). The column temperature was maintained at 40°C, autosampler at 4°C, and injection volume at 3 µL. The mobile phase and gradient were similar as described previously^64^. Mobile phase A: 5 mM ammonium acetate in water (pH=9.0 with addition of ammonium hydroxide), and mobile phase B: 100% acetonitrile. Linear gradient was: 0 min, 85% B; 1.5 min, 85% B; 5.5 min, 35% B; 10.5 min, 35% B; 10.6 min, 10% B; 14 min, 10% B; 14.5 min, 85% B, and 24 min, 85% B. Flow rate was 0.3 mL/min.

The mass spectrometry analysis was performed on an Orbitrap Exploris 480 mass spectrometer, or a Q Exactive-Mass spectrometer (Thermo Scientific) equipped with a heated electrospray ionization (HESI) probe. For polar metabolites, the relevant parameters were listed: heater temperature, 120ᵒC; sheath gas, 30; auxiliary gas, 10; sweep gas, 3; spray voltage, 3.6 kV for positive mode and 2.5 kV for negative mode. The capillary temperature was set at 320°C, and S-lens was 55. The full scan range was set at 60–900 (mass to charge [m/z]). The resolution was set at 70000 (at m/z 200). Customized mass calibration was performed before data acquisition using Xcalibur. The maximum injection time (max IT) was 200 ms. Automated gain control (AGC) was targeted at 3 Å∼ 106 ions. LC-MS peak extraction and integration and metabolic tracing data were analyzed using commercially available Compound Discoverer 3.3 software (Thermo Scientific).

### Statistical Analysis and Bioinformatics

Pathway analysis of metabolites was carried out with MetaboAnalyst 5.0 software (https://www.metaboanalyst.ca/) using the KEGG pathway database (https://www.genome.jp/kegg/). RNA-seq data analysis was mainly performed in R (Version 4.4.2). Differential gene expression analysis was performed with DESeq2, genes were considered significant at |log2FC| > 0.58 and P value <0.05 for RNA-seq data from MC38 and BPD6 tumors, and at |log2FC| > 1 and Padj value <0.05 for RNA-seq data from CD8^+^ T cells. Pathway enrichment of RNA-seq results was carried out using the clusterProfiler package, and gene set enrichment analysis (GSEA) was performed with fgsea package. Additional statistical analysis was performed with GraphPad Prism 10.0 (GraphPad Software), using either a 2-tailed Student’s t test or 1-or 2-way ANOVA. For both 1-way and 2-way ANOVAs, a post-test analysis was performed using Bonferroni’s multiple correction. The number of replicates is indicated in the figure legends. A P value < 0.05 was considered statistically significant.

## Data availability

RNA-seq data have been deposited at the NCBI GEO database upon publication.

## Contribution

X.G., B.C., and C.Y.C. conceived and designed the experiments. X.G., X.Q.and B.C. performed animal experiments. X.G. and X.Q. performed metabolomics analysis. B.C., X.Q., C.Y.C., X.G., S.A., and P. J conducted immunostaining and flow cytometry analysis. X.Q., X.G., P.L., and D.G performed RNA sequencing analysis. Manuscript was written by X.G. with editorial input from B.C., C.Y.C., and K.E.P. X.G. had full access to all the data in the study and takes responsibility for the integrity of the data and the accuracy of the data analysis

## Acknowledgements

This work was funded by NIH/NCI R00 CA237618, USDA 3092-51000-064-05, and the Cancer Prevention and Research Institute of Texas (CPRIT Scholar in Cancer Prevention and Research award RR210056) to X.G. CPRIT Scholar in Cancer Research (RR210029) and V Foundation (V2022-026) to D.G. This project was supported by the Cytometry and Cell Sorting Core at Baylor College of Medicine with funding from the CPRIT Core Facility Support Award (CPRIT-RP180672), the NIH (CA125123 and RR024574) and the assistance of Joel M. Sederstrom. We also thank all the other Gao lab members for their technical assistance and discussion. K.E.P. is an Andrew Sabin Family Foundation Fellow at The University of Texas MD Anderson Cancer Center

## Supplementary Figure Legends

**Fig S1.**
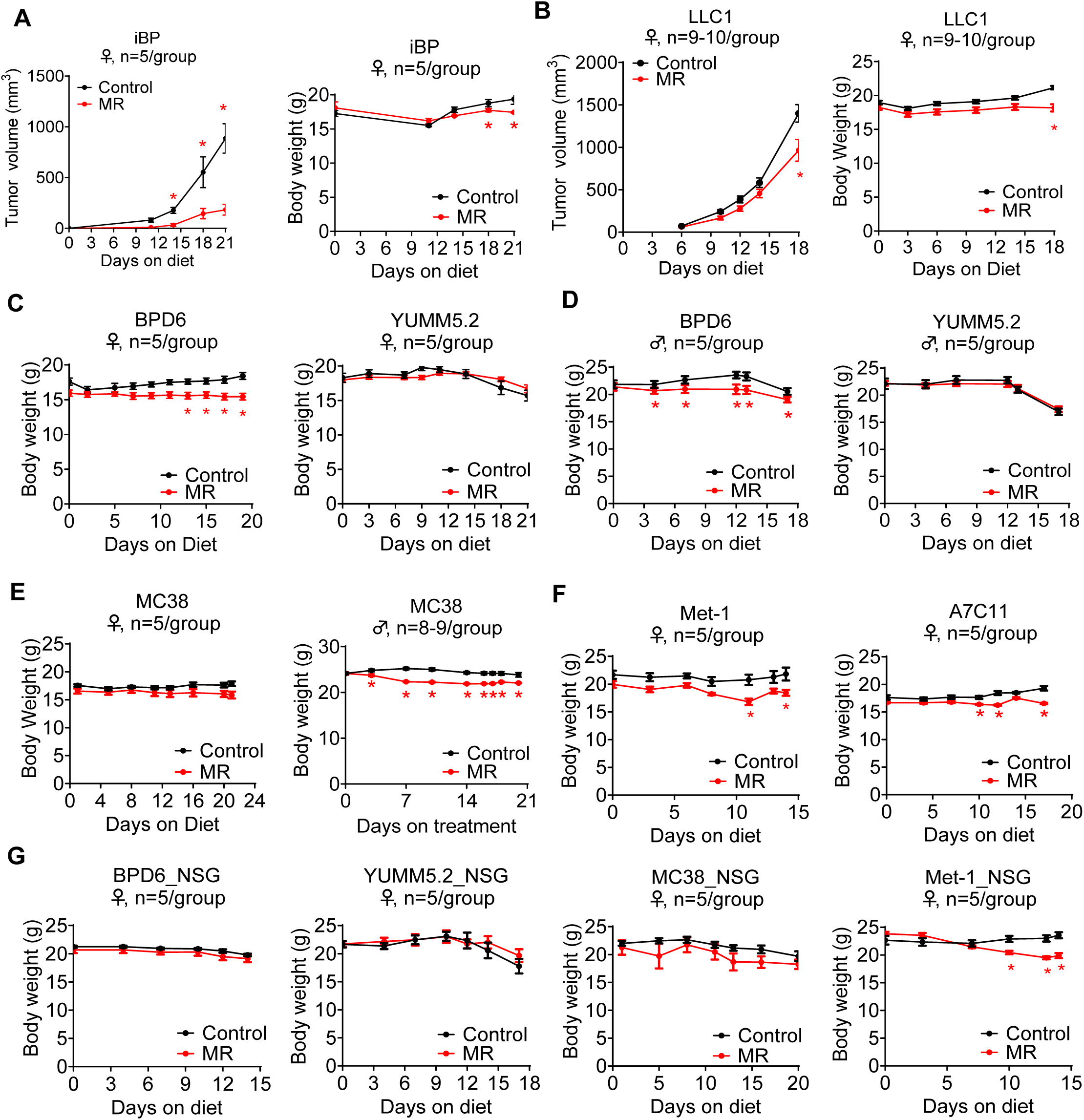
MR inhibits tumor growth in immune-competent mouse cancer models. A. Tumor growth and body weight in female genetic melanoma mouse model iBP. Tumor formation was induced with a single intradermal injection of 150 μg 4-hydroxytamoxifen (*n =* 5/group). B. Tumor growth and body weight in female syngeneic lung cancer model LLC1 (*n =* 9-10/group). C. Body weight in BPD6 and YUMM5.2 syngeneic melanoma models established in female C57BL/6 mice (*n =* 5/group). D. Body weight in BPD6 and YUMM5.2 syngeneic melanoma models established in male C57BL/6 mice (*n =* 5/group). E. Body weight in MC38 syngeneic colorectal cancer model established in female (*n =* 5/group) and male (*n =* 8-9/group) C57BL/6J mice. F. Body weight in Met-1 and A7C11 syngeneic melanoma models established in female FVB and C57BL/6 mice, respectively (*n =* 5/group). G. Body weight in BPD6, YUMM5.2, MC38, and Met-1 models established in female NSG mice (*n =* 5/group). Data were expressed as mean ± SEM. \**P <* 0.05 by 2-way ANOVA followed by Bonferroni’s multiple-correction test.

**Fig S2.**
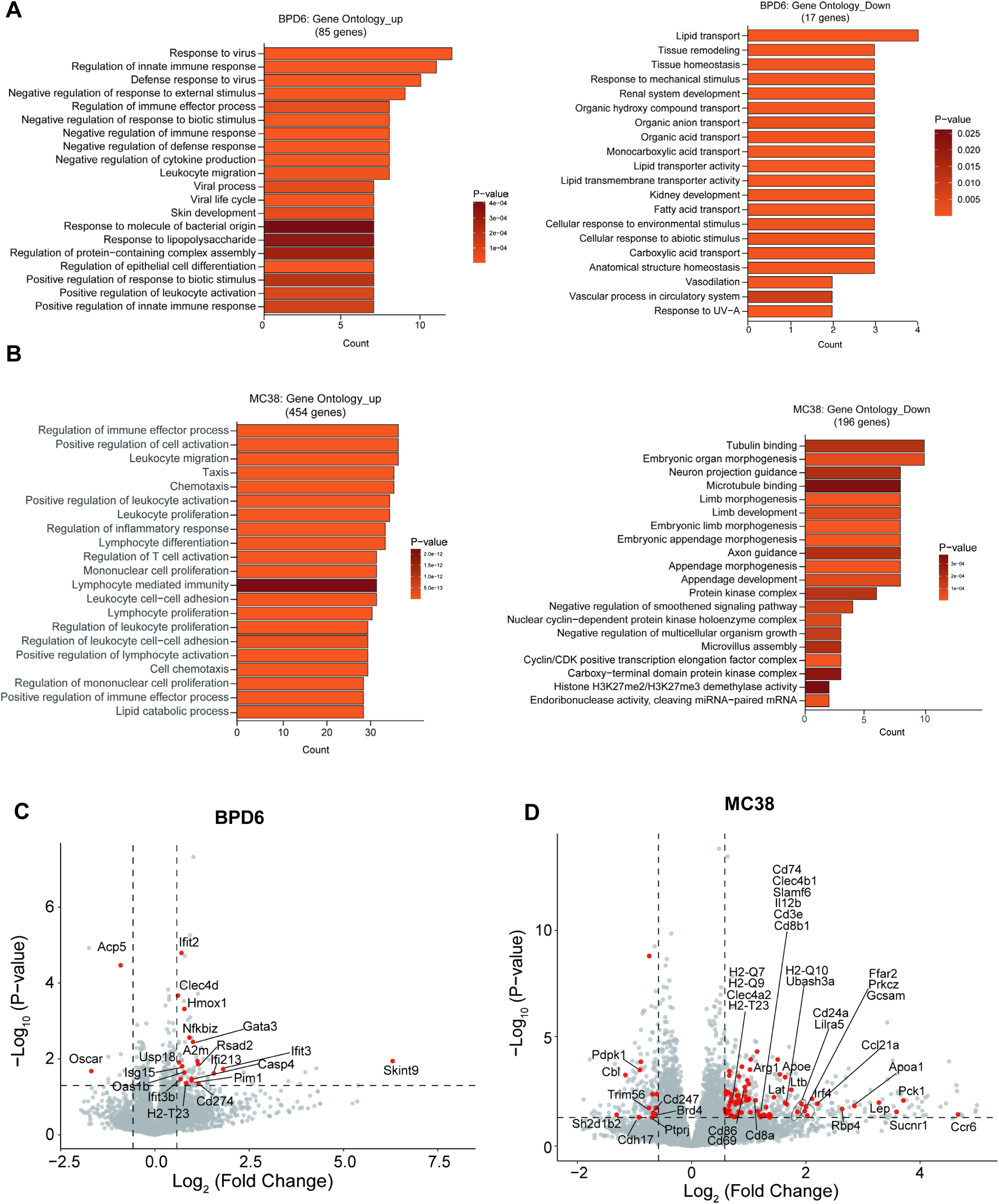
MR boosts intratumor immune response A. The Gene Ontology (GO) process enrichment analysis of 85 significantly up-regulated and 17 significantly down-regulated genes in BPD6 tumors from female C57BL/6 mice under MR vs Control. B. The Gene Ontology (GO) process enrichment analysis of 454 significantly up-regulated and 196 significantly down-regulated genes in MC38 tumors from female C57BL/6 mice under MR vs Control. C. Volcano plot of genes in BPD6 tumors from female C57BL/6 mice under MR vs Control. Colored dots indicated genes related to CD8^+^ T cells D. Volcano plot of genes in MC38 tumors from female C57BL/6 mice under MR vs Control. Colored dots indicated genes related to CD8^+^ T cells

**Fig S3.**
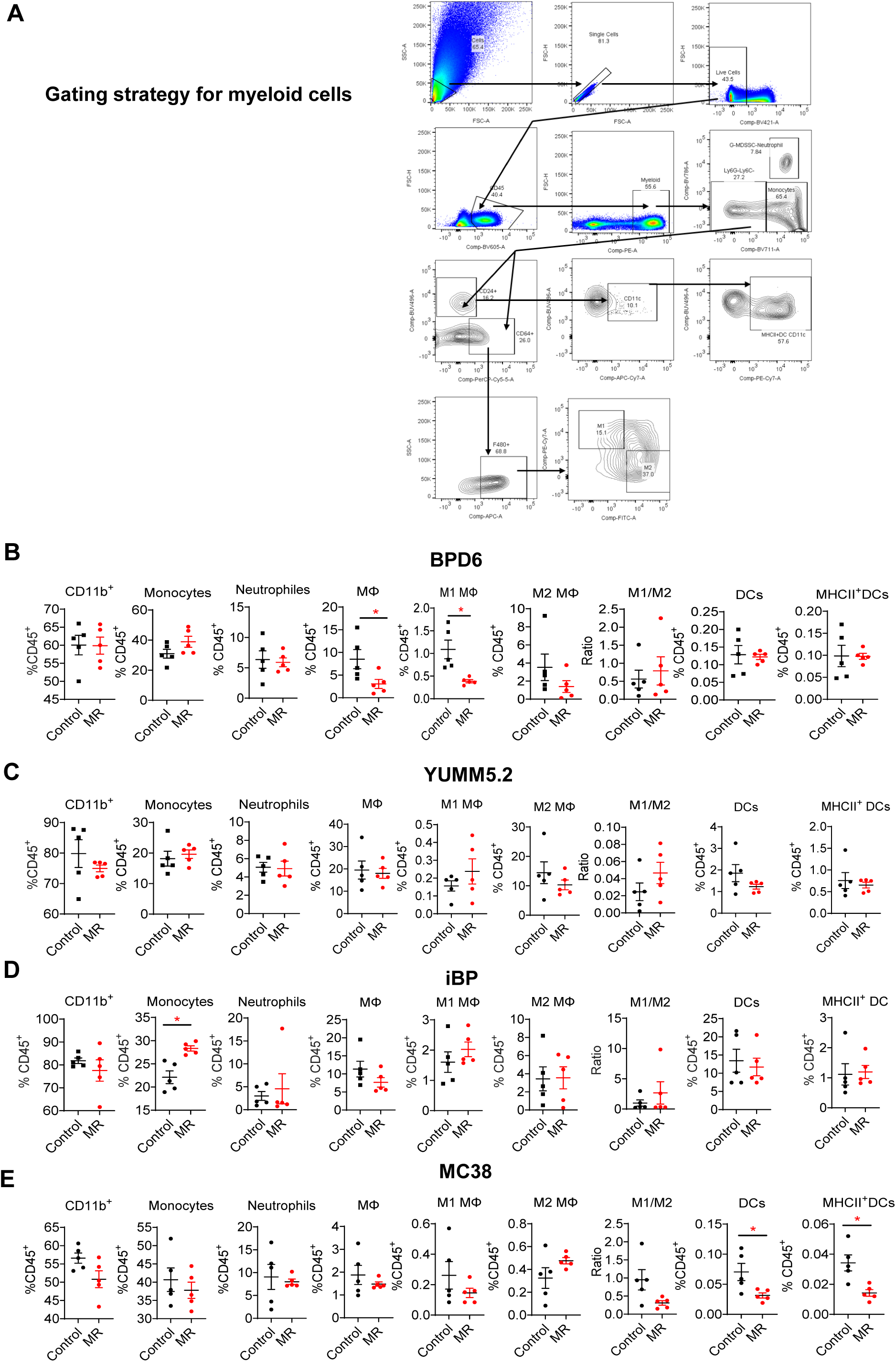
MR shows no consistent effect on intratumor myeloid cells. A. Gating strategy for intratumor myeloid cells by flow cytometry. B. Flow cytometry analysis of myeloid cells in BPD6 tumors in female C57BL/6 mice. C. Flow cytometry analysis of myeloid cells in YUMM5.2 tumors in female C57BL/6 mice. D. Flow cytometry analysis of myeloid cells in iBP tumors in female mice. E. Flow cytometry analysis of myeloid cells in MC38 tumors in female C57BL/6 mice. Data were expressed as mean ± SEM. *N =* 5/group. *P *<* 0.05, by two-tailed Student’s *t* test (B-D). DCs, Dendritic cells; MΦ, macrophages.

**Fig S4.**
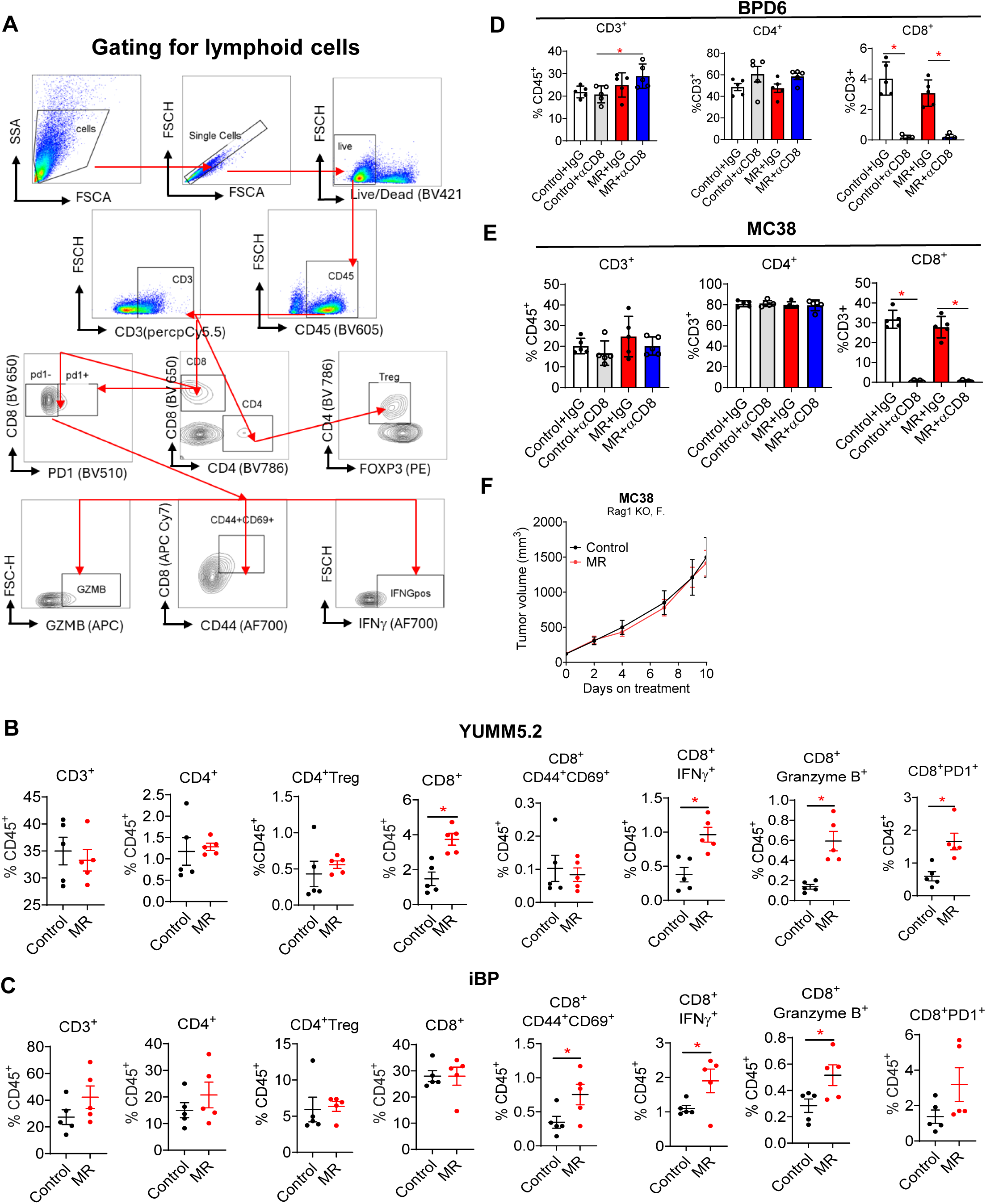
MR inhibits tumor growth via activating tumor infiltrated CD8^+^ T cells A. Gating strategy for intratumor lymphocytes by flow cytometry. B. Flow cytometry analysis of lymphocytes in YUMM5.2 tumors. C. Flow cytometry analysis of lymphocytes in iBP tumors. D. Percentage of CD3^+^, CD4^+^, and CD8^+^ T cells in BPD6 tumors treated with control or MR diet together with αCD8 or IgG isotype treatment. E. Percentage of CD3^+^, CD4^+^, and CD8^+^ T cells in MC38 tumors treated with control or MR diet together with αCD8 or IgG isotype treatment. F. Tumor growth in MC38 and A7C11. Data were expressed as mean ± SEM. *N =* 5/group. *P *<* 0.05, by two-tailed Student’s *t* test (A and B), by 1-way ANOVA (D and E), and by 2-way ANOVA followed by Bonferroni’s multiple-correction test (F).

**Fig S5.**
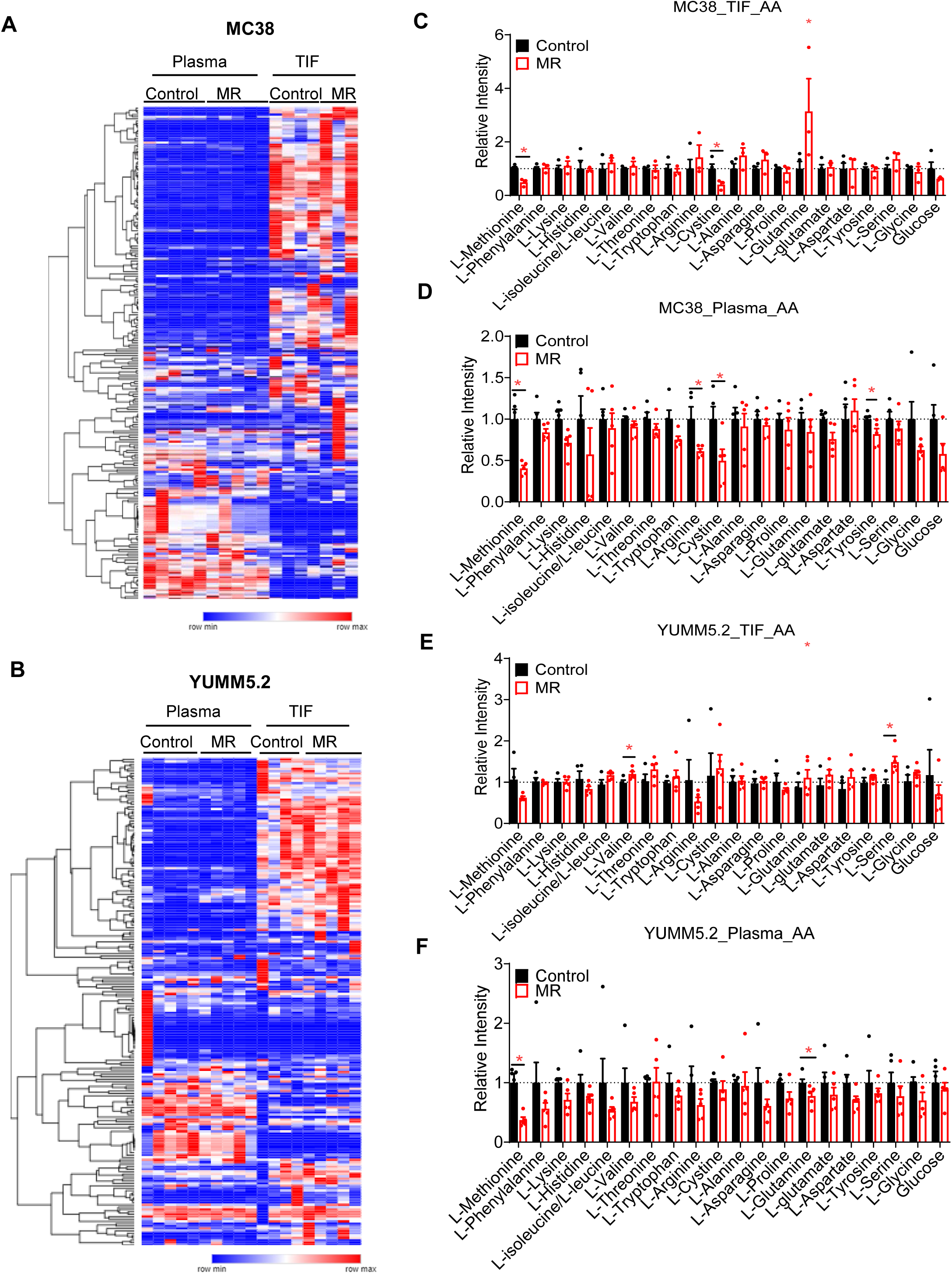
Methionine is not the limiting factor in the tumor microenvironment A. Heatmap of polar metabolites in plasma and TIF from MC38 model. B. Heatmap of polar metabolites in plasma and TIF from Yumm5.2 model. C. Relative mass intensity of amino acids in the TIF from MC38 model. D. Relative mass intensity of amino acids in the plasma from MC38 model. E. Relative mass intensity of amino acids in the TIF from Yumm5.2 model. F. Relative mass intensity of amino acids in the plasma from Yumm5.2 model. Data were expressed as mean ± SEM. *P *<* 0.05 by two-tailed Student’s *t* test (C-F). N*=* 5/group except for MC38-MR-TIF (*n =* 3) and MC38-Control-TIF (*n =* 4).

**Fig S6.**
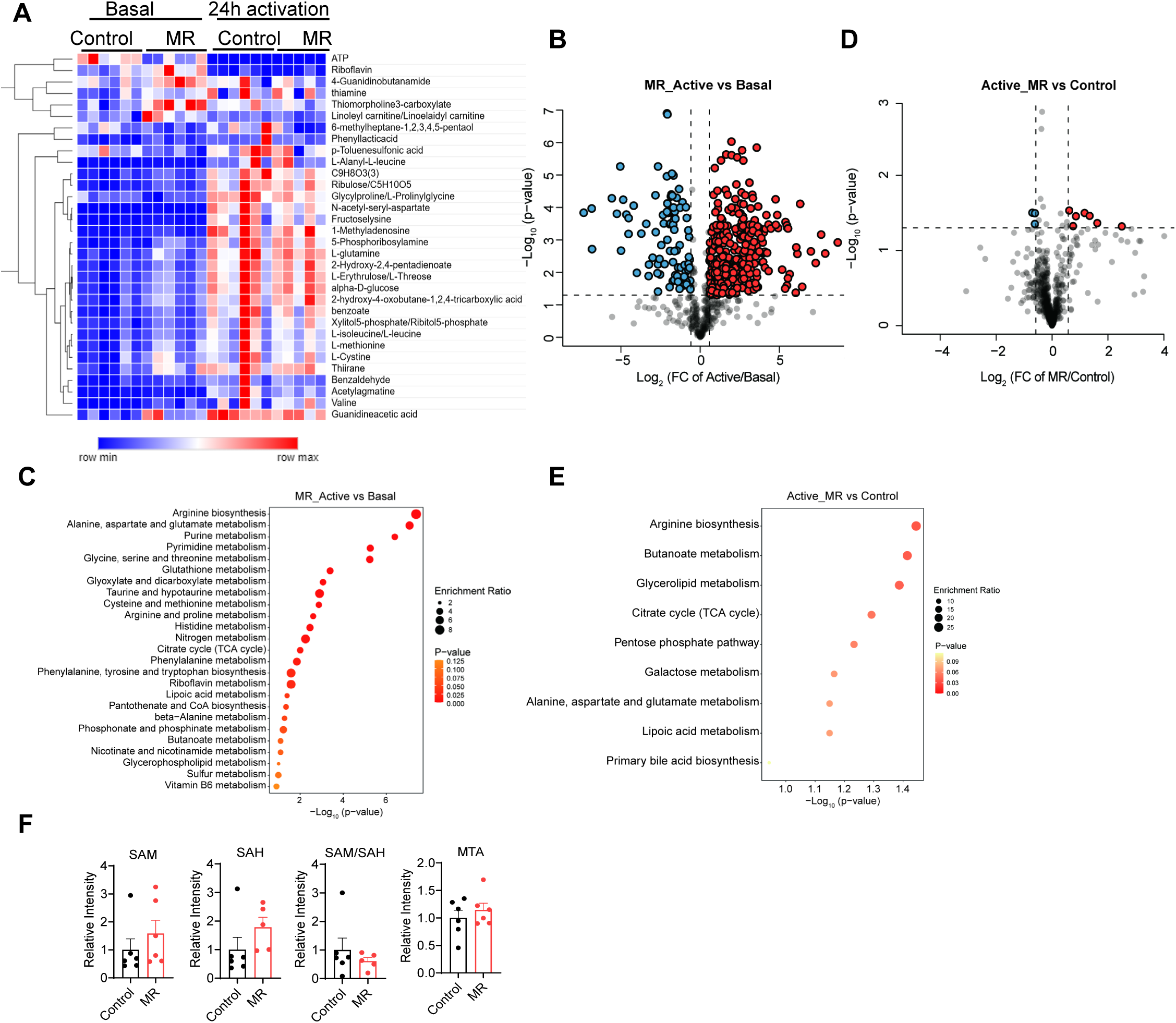
MR primes spleen CD8+ T cell metabolism for activation A. Heatmap of 23 commonly changed metabolites in Figure 4D and 4F (*n =* 5-6/group). B. Volcano plot of 684 metabolites in CD8+ T cells isolated from mice fed MR diet before and after activation. Colored dot indicates that P < 0.05 & |log_2_FC| > 0.58. C. Gene Ontology analysis of significantly changed (*P* < 0.05 & |log_2_FC| > 0.58) genes by MR in B. D. Volcano plot of 684 metabolites in CD8+ T cells isolated from mice fed control or MR diet after activation. Colored dot indicates that P < 0.05 & |log_2_FC| > 0.58. E. Gene Ontology analysis of significantly changed (*P* < 0.05 & |log_2_FC| > 0.58) genes by MR in D. F. Relative intensity of SAM, SAH, MTA (5-methylthioadenosine), and relative ratio of SAM/SAH in CD8+ T cells isolated from mice fed control or MR diet under basal condition before activation. Data were expressed as mean ± SEM. *P *<* 0.05 by two-tailed Student’s *t* test.

**Fig S7.**
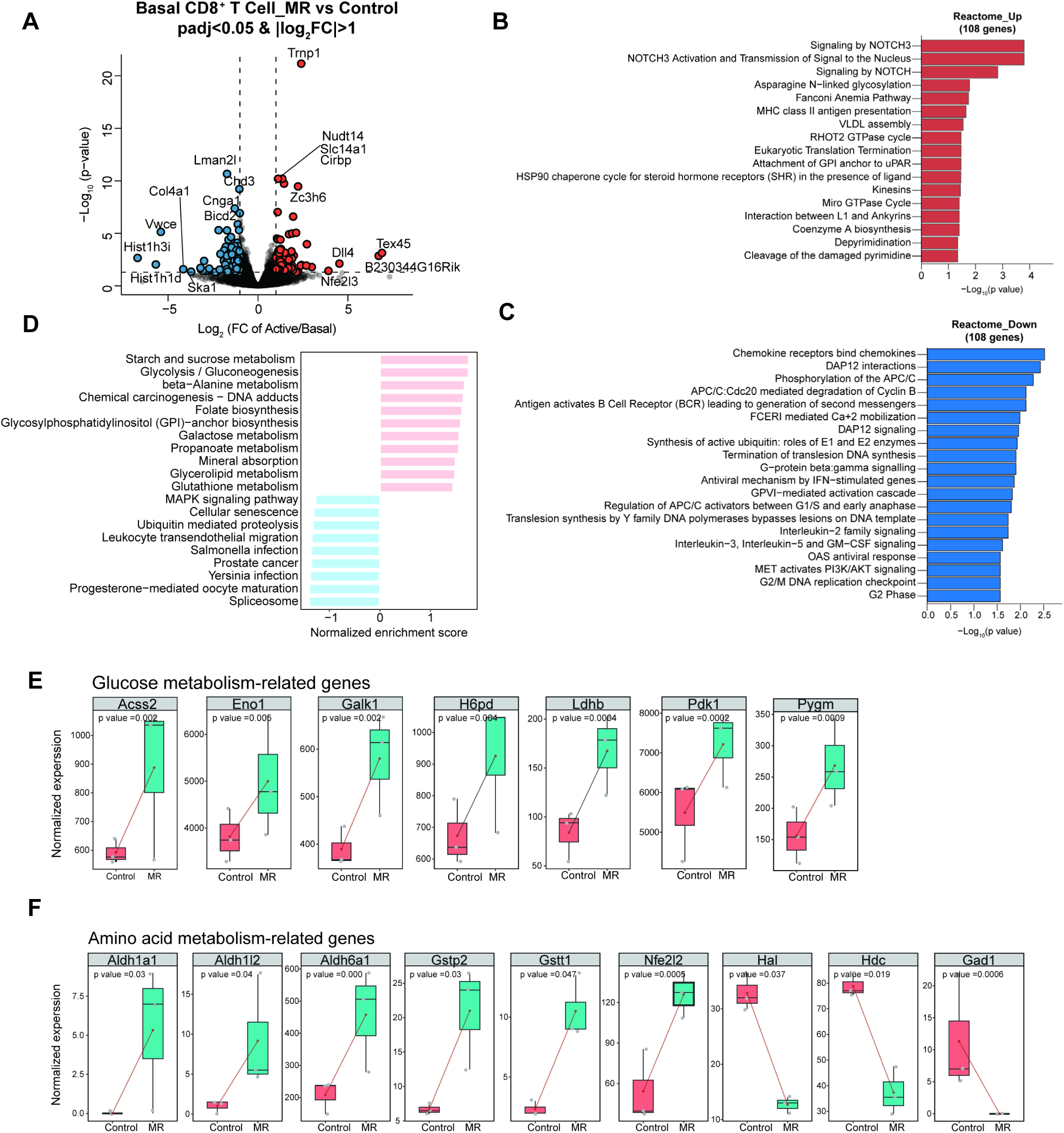
MR affects gene expression in spleen CD8+ T cell A. Volcano plot of genes changed in CD8+ T cells by MR under basal condition (n = 3/group). Red (up-regulated) and blue (down-regulated) dots indicate that Padj < 0.05 & |log_2_FC| > 1. B. Gene Ontology analysis of 108 significantly up-regulated Padj < 0.05 & |log_2_FC| > 1 genes by MR in A. C. Gene Ontology analysis of 108 significantly down-upregulated Padj < 0.05 & |log_2_FC| > 1 genes by MR in A. D. Pathway alternation of CD8^+^ T cells from control or MR-fed mice before activation (n = 3/group) was evaluated using GSEA. E. Boxplot of top differentially expressed genes in glucose metabolism-related pathways. F. Boxplot of top differentially expressed genes in amino acid metabolism-related pathways. Data were expressed as mean ± SEM. *P *<* 0.05 by two-tailed Student’s *t* test (E-F).

**Fig S8.**
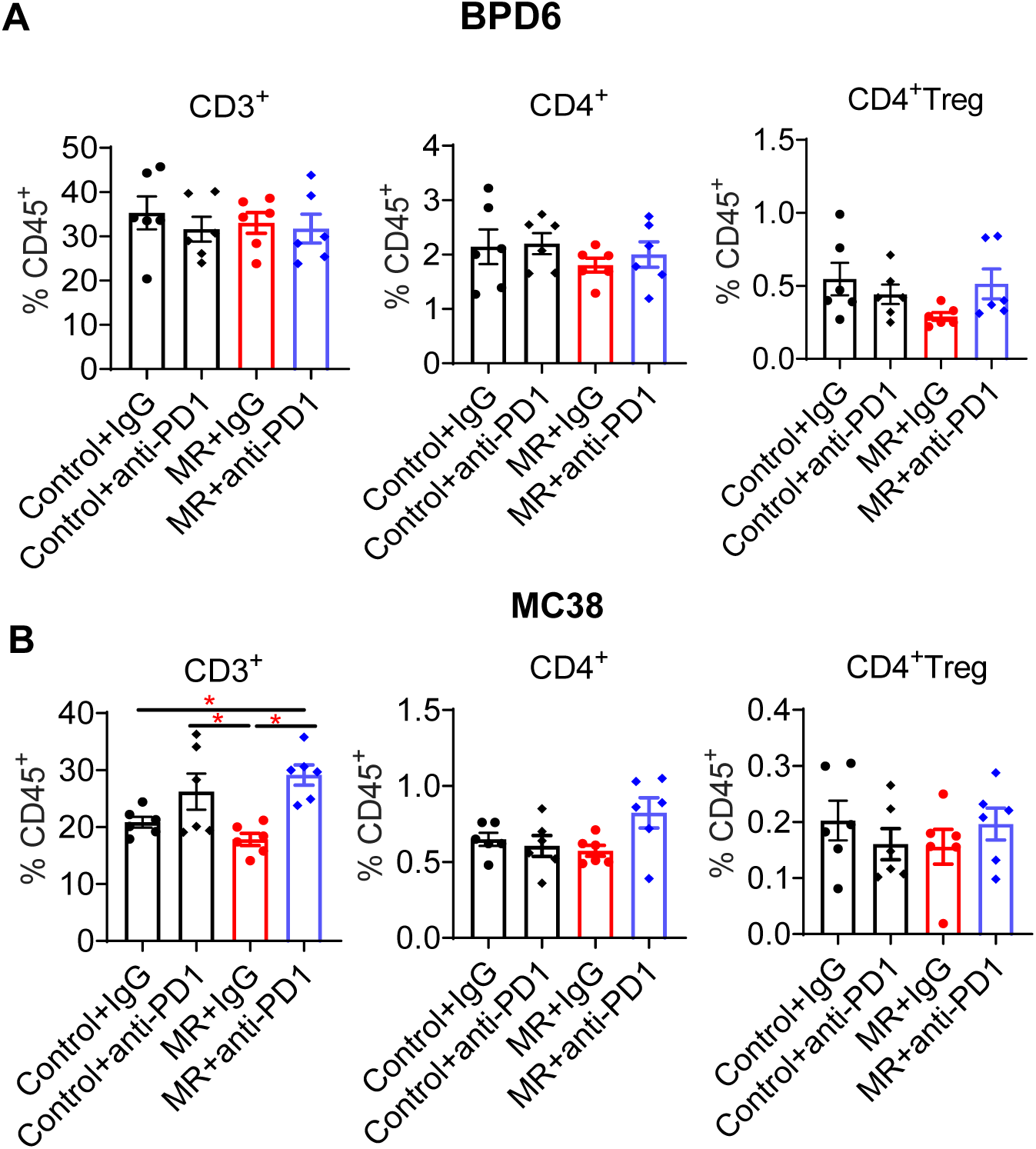
MR synergizes with anti-PD1 treatment. A. Flow cytometry analysis of intratumor T cell markers in BPD6 tumors: CD3^+^, CD4^+^, and CD4^+^Treg (CD4^+^Foxp^+^) T cells (*n =* 5/group). B. Flow cytometry analysis of intratumor T cell markers in MC38 tumors: CD3^+^, CD4^+^, and CD4^+^Treg (CD4^+^Foxp^+^) T cells (*n =* 5/group). Data were expressed as mean ± SEM. *P < 0.05 by one-way ANOVA test unless otherwise indicated.

